# Mechanoreceptor synapses in the brainstem shape the central representation of touch

**DOI:** 10.1101/2021.02.02.429463

**Authors:** Brendan P. Lehnert, Celine Santiago, Erica L. Huey, Alan J. Emanuel, Sophia Renauld, Nusrat Africawala, Ilayda Alkislar, Yang Zheng, Ling Bai, Charalampia Koutsioumpa, Jennifer T. Hong, Alexandra R. Magee, Christopher D. Harvey, David D. Ginty

**Author notes:** Co-first author (Brendan P. Lehnert, Celine Santiago).

## Abstract

Mammals use glabrous (hairless) skin of their hands and feet to navigate and manipulate their environment. Cortical maps of the body surface across species contain disproportionately large numbers of neurons dedicated to glabrous skin sensation, potentially reflecting a higher density of mechanoreceptors that innervate these skin regions. Here, we find that disproportionate representation of glabrous skin emerges over postnatal development at the first synapse between peripheral mechanoreceptors and their central targets in the brainstem. Mechanoreceptor synapses undergo developmental refinement that depends on proximity of their terminals to glabrous skin, such that those innervating glabrous skin make synaptic connections that expand their central representation. In mice that do not sense gentle touch, mechanoreceptors innervating glabrous skin still make more powerful synaptic connections in the brainstem. We propose that the skin region a mechanoreceptor innervates controls refinement of its central synapses over development to shape the representation of touch in the brain.

## Introduction

Perception relies on populations of sensory neurons that represent salient features of the physical world. Neurons are not uniformly allocated across a sensory space, but rather, particular regions are often represented by disproportionate numbers of neurons. A striking example is the primate visual system, where neurons across the visual hierarchy disproportionately represent central regions of the visual field, in part reflecting the high density of photoreceptors in the center of the primate retina (Cowey and Rolls, 1974). Disproportionate central representation of foveal visual space has far-reaching consequences, as this is thought to underlie ocular motor strategies used to survey the world as well as cognitive strategies used to comprehend visual scenes (Yarbus, 1967). Similarly, the human somatosensory cortex contains an iconic map of the body, the homunculus, that does not faithfully reflect our physical form, but rather disproportionately allocates neurons to the hands and face at the expense of the torso and proximal limbs (Penfield and Boldrey, 1937). Disproportionate central representation may reflect the tension between capturing the entirety of a sensory space and preferentially allocating limited neural resources to areas where acuity or representational capacity is critical.

In mammals, skin can be broadly classified as hairy or glabrous (non-hairy), and a common feature of mammalian cortical body maps is that they strongly favor glabrous skin surfaces (Penfield and Boldrey, 1937; Sur et al., 1978; Sur et al., 1980; Xu and Wall, 1999). In primates, for example, glabrous skin representations occupy 30% of somatosensory cortex but only 5% of skin surface area (Sur et al., 1982). Glabrous skin found on distal limbs is molecularly and structurally distinct from the hairy skin covering the majority of the body, and it contains specialized corpuscle-associated sensory endings important for object manipulation and touch discrimination (Abraira and Ginty, 2013; Lu et al., 2016). Touch-sensitive neurons more densely innervate skin areas with high tactile acuity, such as the palm of the hands, and these peripheral innervation density gradients are purported to underlie the distorted figure of the cortical somatosensory homunculus (Catani, 2017). This view is supported by observations in rodents, where neurons in layer IV somatosensory cortex that respond to whisker stimulation are organized into barrels, and anatomical measurements revealed that the number of neurons in each barrel is proportionate to the number of myelinated sensory neurons innervating the corresponding whisker (Lee and Woolsey, 1975).

However, multiple lines of evidence challenge the conventional view that disproportionate central representation of the body is simply a reflection of skin innervation density. For example, the primary somatosensory cortex and the superior colliculus contain body maps with differing proportions (Drager and Hubel, 1976), suggesting that the same peripheral inputs can support different central representations. Moreover, a recent compilation of human anatomical and physiological measurements argues that differences in peripheral sensory neuron numbers may not account for the over-representation of the hands and the digits in somatosensory cortex (Corniani and Saal, 2020). This parallels work in the visual system, where studies argue that increased photoreceptor density at the primate retinal fovea alone is insufficient to explain the enlarged representation of central fields of view in the visual cortex (Azzopardi and Cowey, 1993). Thus, despite the prevalence of disproportionate body maps across mammals with diverse body forms, the mechanisms that shape the representation of the body in the brain have remained incompletely understood since the discovery of these maps nearly a century ago.

Here, we examine the central representation of glabrous and hairy skin in mice and find that disproportionate emphasis of glabrous skin emerges during postnatal development. At early postnatal stages, the number of neurons in the mouse brain responsive to a body region reflects the density of receptors in the skin. Later, developmental refinement of synaptic connections in the brainstem produces a synaptic expansion that favors glabrous skin surfaces in adulthood. Sparse peripheral mechanoreceptor labeling experiments reveal that the structural complexity and numbers of synapses made by mechanoreceptors depend on the skin region they innervate, with glabrous skin-innervating neurons exhibiting more complex arbors and synapses in the brainstem. Moreover, optical stimulation of glabrous skin innervating mechanoreceptors evokes larger synaptic currents and is more effective at driving spiking in brainstem neurons than stimulation of hairy skin innervating sensory neurons. Interestingly, this difference is preserved in mice that do not sense gentle touch on their body surface. Together, our findings support a model in which peripheral innervation density and skin region dependent synaptic expansion in the brainstem act multiplicatively to shape the central representation of touch.

## Results

### Disproportionate expansion of glabrous skin representation emerges over postnatal development while receptor density in the skin remains stable

We first mapped the extent of hindlimb and forelimb primary somatosensory cortex using multiphoton imaging in adult mice (Figure 1A). A greater proportion of Layer IV neurons in primary somatosensory cortex (S1) responded preferentially to stroking across glabrous skin of the paw compared to stroking hairy skin of the paw, with glabrous skin and hairy skin S1 neurons displaying an intermingled, salt-and-pepper organization. Recordings from low-threshold mechanoreceptors (LTMRs) in the L4 dorsal root ganglion (DRG) confirmed that the light stroke stimulus used for these functional mapping experiments is an equally effective stimulus for LTMRs that innervate glabrous or hairy skin of the paws (Supplemental Figure 1). To characterize emerging body representations over development, hindlimb and forelimb S1 were targeted with tiled multielectrode array penetrations at 100-200 micron intervals in postnatal day 14 mice, an early developmental period that coincides with eye opening and precedes weaning. Recordings from hindpaw regions of P14 cortex showed an equal number of S1 neurons responding to paw glabrous and paw hairy skin stimulation, while forepaw cortex more strongly emphasized paw glabrous skin surfaces (Figure 1B, C). Similar recordings in adult mice uncovered a dramatically more pronounced glabrous representation bias, an approximately 3-fold increase in paw glabrous-to-hairy skin preference across both hindpaw and forepaw S1, compared to P14 animals (Figure 1B, C).

**Figure 1.**
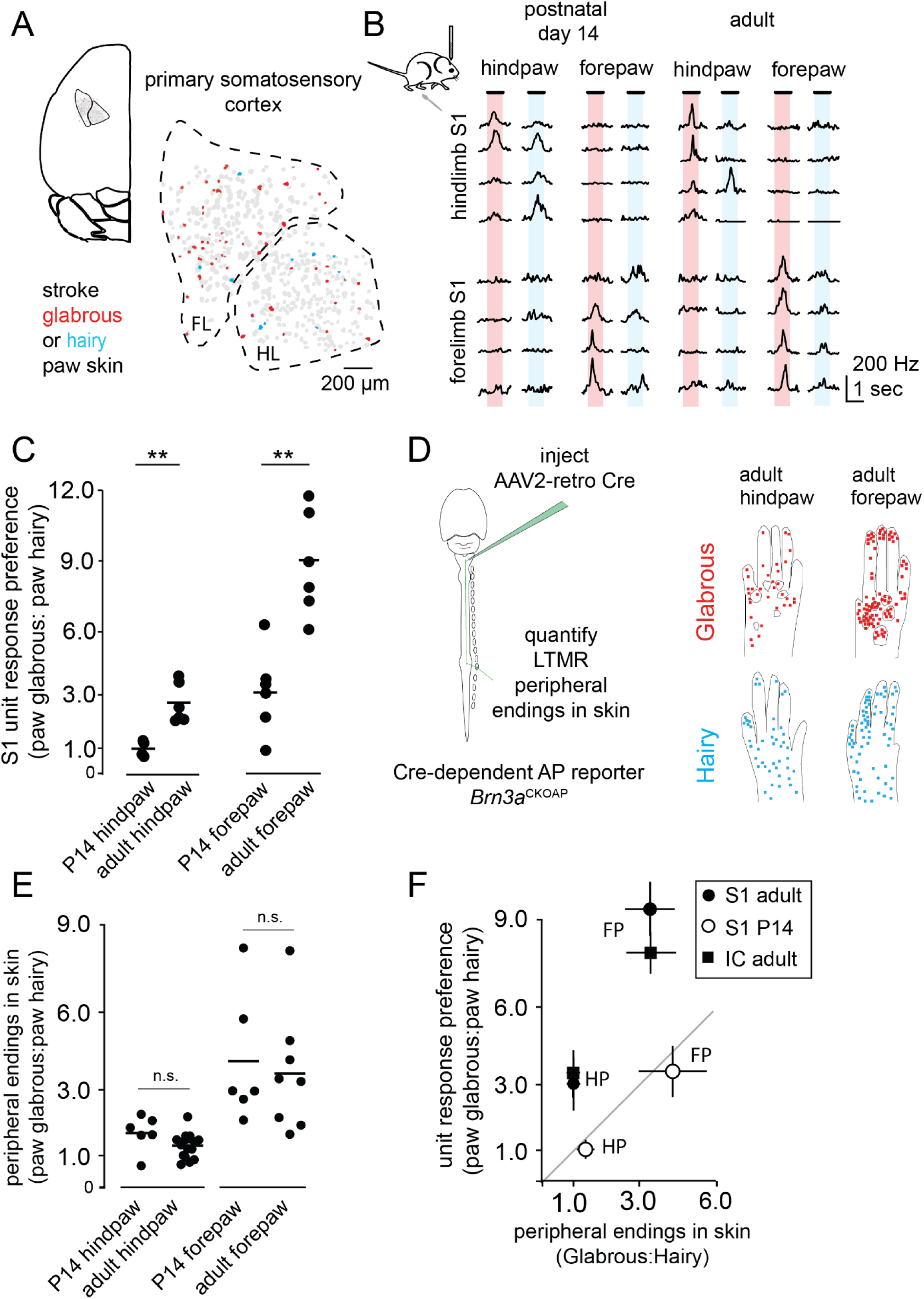
Central representations of glabrous skin expand over postnatal development while receptor density is stable. **A**. Preference of cells in layer IV mouse somatosensory cortex (S1) for light tactile stimulation of paw glabrous (red) or paw hairy (blue) skin. Multiphoton functional maps were used to guide multielectrode array (MEA) recordings of S1 in adult and postnatal day 14 (P14 mice) with ~200 μm spacing between penetrations. **B**. MEA recordings from S1 at two developmental time points. Representative trial-averaged responses to touch of glabrous and hairy skin for hindpaw (top) and forepaw (bottom) regions of somatosensory cortex reveal a preference for glabrous skin stimulation that emerges over development. **C**. The ratio of neurons preferentially responding to gentle stroke of glabrous or hairy skin for forepaw S1 and hindpaw S1. An increase in the proportion of neurons preferring glabrous skin stroke occurs over development (p < 0.01, Mann-Whitney U test). **D**. (left) Strategy for estimating innervation density across the body. AAV2-retro-Cre virus was injected into the dorsal column of *Brn3AP_CKOAP_* reporter mice at concentrations that label countable numbers of Aβ LTMR endings in the skin. (right) Compilation of all endings identified in the paws of adult animals that received dorsal column injections, super-imposed on reference images of paws. Each dot represents a single Aβ LTMR terminal ending in skin. **E**. The relative number of Aβ LTMR endings in glabrous or hairy forepaw or hindpaw skin is stable from P14 into adulthood (Welch’s t-test, n=6 animals for P14, n=12 animals for adult hindpaw skin, n=7 animals for adult forepaw skin). Each dot on the plot indicates a distinct paw; both left and right paws were analyzed for some animals. For each paw, glabrous:hairy innervation density ratios were calculated by dividing the number of endings per square mm in paw glabrous skin by the number of endings per square mm in paw hairy skin. **F**. (left) The relationship between innervation density and the proportion of neurons preferring glabrous skin touch in juvenile and adult mice. At P14, glabrous skin preference ratios for both forepaw (FP) S1 and hindpaw (HP) S1 are proportionate to innervation density. In adults, a greater proportion of S1 neurons respond to glabrous skin stimulation than predicted by innervation density, and a similar effect is seen in the inferior colliculus, a second major tactile area (right). Together, this demonstrates a developmental expansion of glabrous skin representation in multiple brain regions that is independent of receptor density.

Sensory neuron innervation density differences across skin regions are thought to account for the disproportionate central representation of touch. In the visual system, the heightened photoreceptor density characteristic of the primate fovea emerges over weeks of postnatal development (Packer et al., 1990), and so we next asked whether a similar developmental rearrangement of peripheral touch neurons accompanies the dramatic developmental shift in paw glabrous skin and hairy skin functional representation observed in S1. Touch is transduced by a heterogeneous population of LTMRs, which extend an axonal branch with highly specialized mechanosensitive endings to the skin, and functional responses in primate S1 depend on Aβ-LTMRs that ascend the dorsal column of the spinal cord and project directly to the brainstem (Jain et al., 1997). We measured the distribution of the terminal endings of these neurons in the skin through retrograde viral transduction of sensory neurons that ascend the dorsal column, using the Cre-dependent alkaline phosphatase (AP) reporter *Brn3a^cKOAP^* (Figure 1D)(Badea et al., 2009b). Viral concentrations were titrated such that neurons were sparsely labeled and the endings of individual neurons in the skin were distinguishable (see Methods); all known Aβ-LTMR subtypes were detected in these experiments (Wu et al., 2012; Bai et al., 2015).

We collected the skin of animals at P14 or ~P50 (adult) and performed whole-mount alkaline phosphatase (AP) staining to quantify the relative density of peripheral arbors in each skin area (Figure 1D). We detected stark differences in innervation density along the proximal-distal axis of the limbs of adult mice, consistent with previous electrophysiological and anatomical measurements in other species (Wang et al., 1997) (Supplemental Figure 1). In addition, we found that at both P14 and adulthood, forepaw glabrous skin is approximately three times more densely innervated by Aβ-LTMRs than forepaw hairy skin (Figure 1E, F). In contrast, hindpaw glabrous and hindpaw hairy skin have equal innervation densities at both ages (Figure 1E, F). For each paw, the ratio of innervation densities is expressed as a glabrous: hairy ratio (Figure 1E, F). Between P14 and adulthood, we detect no significant changes in glabrous: hairy skin Aβ-LTMR innervation ratios in either the forepaws or the hindpaws (Figure 1E). Because changes in skin area coverage could occur independently of changes in neuron numbers, for example if peripheral arbors undergo structural remodeling during postnatal development, we also measured the peripheral arbor morphologies of sparsely labeled neurons in glabrous and hairy skin and found no changes in surface area or number of end points between P14 and adulthood (Supplemental Figure 1).

These functional and anatomical findings define the relationship between paw glabrous and paw hairy skin innervation density and central (S1) representation over development. At P14, paw glabrous skin and hairy skin representation in S1 is proportionate to skin innervation density (Figure 1F). In contrast, in adults, there is an approximately 3-fold expansion of glabrous skin preferring S1 neurons beyond that predicted by the distribution of peripheral mechanoreceptors. To examine the proportionate representation of the body in the brain more generally, we next extended our physiological recordings to the external nucleus of the inferior colliculus, another major tactile area within the brain that exhibits a prominent representation of the body surface (Figure 1F, data not shown). As in S1, representations of forepaw and hindpaw glabrous skin and hairy skin regions in the inferior colliculus of adult mice far exceed that predicted by the relative numbers of LTMR endings in these skin regions. Taken together, these findings demonstrate that the degree of central representation of paw glabrous skin and paw hairy skin surfaces is consistent with receptor density early in postnatal development, but that there is a dramatic and unexplained expansion of cortical and collicular neurons responding to touch of glabrous skin in adulthood.

### *In vivo* multiphoton imaging of neurons in the gracile nucleus of the brainstem reveals a cellular locus of expansion

Developmental expansion of glabrous skin preferring neurons in cortex (S1) and inferior colliculus might reflect expansion at a common locus of ascending LTMR inputs within the brainstem. Therefore, we performed dual retrograde labeling experiments targeting the ventroposteriolateral (VPL) thalamus (the main input to somatosensory cortex) and the inferior colliculus to visualize cell bodies of ascending projection neurons of the brainstem; these labeling experiments confirmed dense populations of VPL and inferior colliculus projection neurons within the gracile nucleus (GN) and cuneate nucleus (CN) of the brainstem dorsal column nuclei, a region that receives direct synaptic input from Aβ-LTMRs. Indeed, this dual-color retrograde labeling in a *Gad2^nls-mcherry^* mouse demonstrated the presence of substantial, largely non-overlapping and intermingled populations of thalamic projection neurons, inferior collicular projection neurons, and presumptive local inhibitory neurons, within both the GN and CN (Supplemental Figure 2).

We used this retrograde labeling strategy to selectively express the calcium indicator jGCaMP7s in GN/CN projection neurons and developed a preparation that provides *in vivo* optical access to the GN. The GN, which encodes mechanosensory inputs from the caudal half of the animal, including the hindlimbs, is optically accessible in acute preparations that expose the brainstem, while the CN lies beneath ascending fiber tracts. This imaging technique allows observation of the tactile responses of hundreds of neurons at depths up to ~250 μm and throughout the entirety of the GN for approximately six hours (Figure 2A, data not shown).

**Figure 2.**
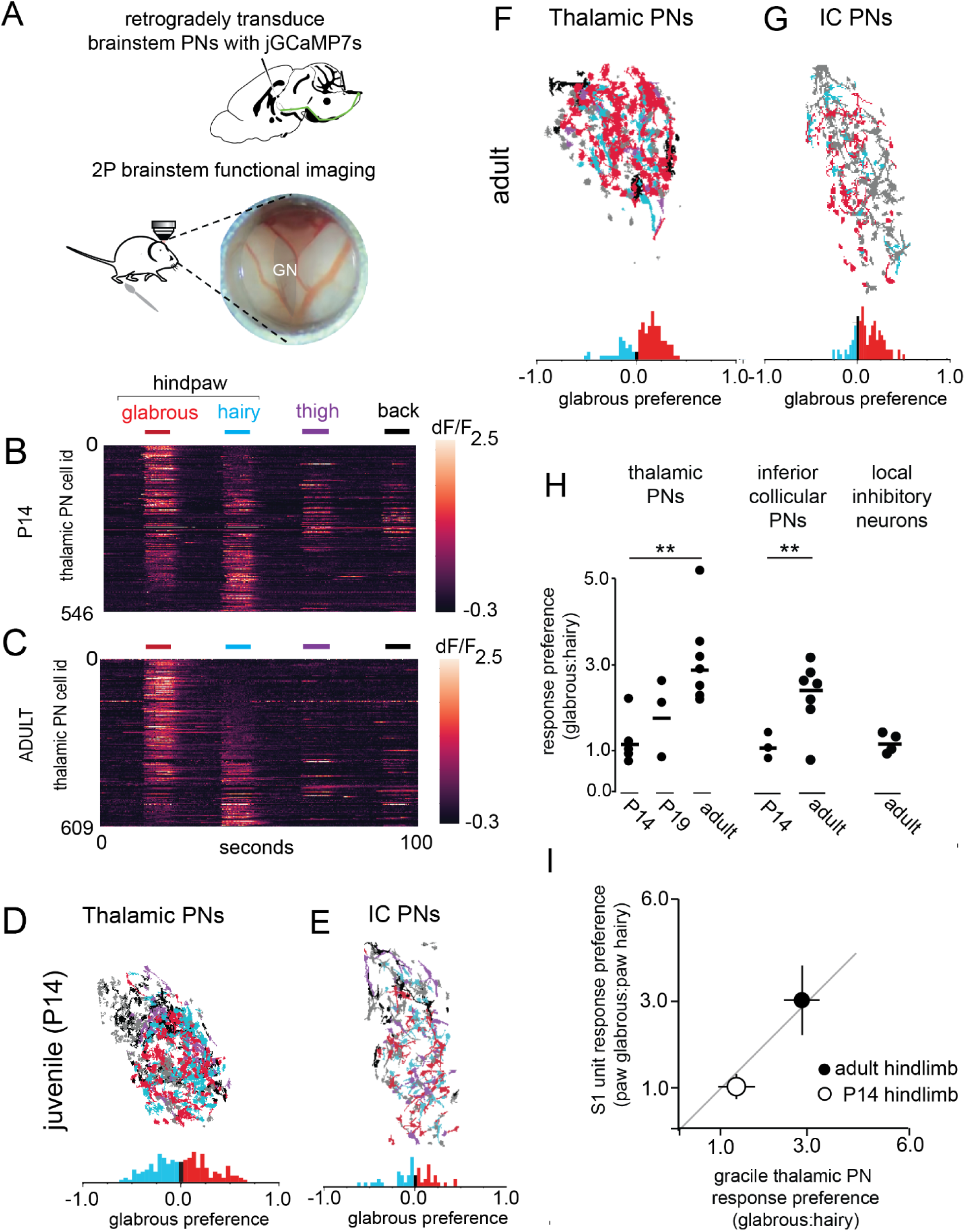
Disproportionate representation of glabrous skin emerges in the brainstem over postnatal development. **A**. (top) Thalamic projecting neurons in the gracile and cuneate nuclei of the brainstem express the calcium indicator jGCaMP7s after retrograde viral transduction of their axonal projections. (bottom) A preparation allowing i*n vivo* multiphoton imaging of the gracile nucleus of the brainstem. **B**. Touch-evoked responses of thalamic projection neurons functionally imaged in the gracile nucleus. Calcium signals were evoked by stroking equivalent skin areas of the hindpaw glabrous skin (red), hindpaw hairy skin (blue), thigh (purple), and back (black) in P14 mice. **C**. Representative response to touch of the body in adult mice, similar to B. **D**. Representative map of the body within gracile nucleus thalamic projection neurons. A glabrous skin preference index, (glabrous – hairy) / (glabrous + hairy), was computed for all neurons responding to hindpaw skin stimulation in this and other experiments (bottom). At P14, there are roughly equal proportions of glabrous preferring and hairy preferring cells. **E**. Representative map of the body within gracile nucleus inferior collicular projection neurons, similar to D. **F-G.** Representative maps of the body in adulthood for thalamic projecting neurons (F) and inferior collicular projecting neurons (G). **H**. The proportion of thalamic or inferior collicular projecting neurons in the gracile nucleus that preferentially respond to hindpaw glabrous skin touch expands over developmental time (p < 0.01, Mann Whitney U test). **I**. The relationship between glabrous preference in the gracile nucleus of the brainstem and glabrous preference in hindpaw S1 at two developmental timepoints.

The GN functional imaging preparation was used to ask whether disproportionate representation of glabrous skin is present within the GN and if so, whether it similarly emerges over postnatal development. Stroke was first delivered to the skin of P14 mice over approximately 10 mm^2^ of the ipsilateral hindpaw glabrous skin, hindpaw hairy skin, thigh, and back. GN neurons responded robustly and repeatedly to tactile stimulation of the ipsilateral lower body, including the ipsilateral hindlimb and hindpaw, while tactile stimulation of the contralateral lower body failed to activate GN neurons (Figure 2B, data not shown). Spatial maps of the body revealed a core region of intermingled hindpaw glabrous skin and hairy skin preferring neurons and less numerous populations of thigh and back preferring neurons. At P14, GN thalamic projection neurons showed no preference at the population level for touch to glabrous skin compared to hairy skin of the hindpaw (Figure 2B, D). The same was true for projection neurons to the inferior colliculus, which exhibited no population bias towards glabrous skin at P14 (Figure 2E).

Strikingly, in adult mice we observed an approximate 3-fold increase in the number of GN thalamic projection neurons preferentially responsive to glabrous skin over those that responded to touch on a comparable surface area on the hairy skin side of the same paw (Figure 2F, H). GN inferior collicular projecting neurons also showed a marked preference for glabrous skin stimulation (Figure 2G,H), while expanded glabrous skin representations were not observed for locally projecting inhibitory neurons (Figure 2H). It is noteworthy that GN thalamic projection neuron populations had glabrous skin to hairy skin preference ratios that were equivalent to cortical (S1) neuron preference ratios measured at both P14 and adult (Figure 2I, Mann-Whitney U test, all p > 0.05). Taken together, these findings indicate that disproportionate representation of glabrous skin in both S1 and the inferior colliculus is partially or fully driven by changes in the GN over development. As Aβ-LTMR innervation density is stable over this time period, we hypothesized that disproportionate representation of glabrous skin in the brainstem emerges via developmental changes in synaptic connectivity. A testable prediction of this hypothesis is that the number or function of LTMR synapses in the brainstem is a reflection of the skin type innervated.

### Differences in the morphology of LTMR axonal central projections and the number of LTMR synapses in the brainstem reflect innervating skin type and developmental stage

Aβ-LTMRs can project nearly the length of the entire body, from peripheral skin mechanosensory terminals in the distal limbs to their central synaptic terminals in the GN and CN of the brainstem. We sparsely labeled Aβ-LTMRs to explore the relationship between the skin type innervated and the anatomy of their axonal arbors in the GN and CN. The three main classes of cutaneous Aβ-LTMRs were stochastically labeled using the Cre-dependent AP reporter mouse *Brn3a^CKOAP^* in conjunction with LTMR-selective CreER lines and low-dose tamoxifen treatments (see Methods). Samples had 1- 4 labeled LTMRs per animal with central terminals projecting to the GN or CN, such that the peripheral arbors of individual LTMRs could be identified in the skin based on established somatotopy. This approach was effective in generating sparsely labeled trunk-level Aβ-LTMRs, but more rarely resulted in paw-innervating LTMR labeling. Thus, we generated a dual Cre and FlpO recombinase-dependent *Tau^FsF-iAP^* reporter mouse to intersectionally label neurons innervating paw glabrous or paw hairy skin through injection of an AAV2/1-FlpO virus to the skin of animals containing CreER alleles (Figure 3A and Supplemental Figure 3). We used *Ret^CreER^, TrkC^CreER^*, or *Ptgfr^CreER^* to label Aβ rapidly adapting (RA)-LTMRs in hairy and glabrous skin (*Ret^CreER^*); Aβ slowly adapting (SA) type I-LTMRs in hairy and glabrous skin (*TrkC^CreER^*), or Aβ Field-LTMRs in hairy skin and a previously uncharacterized Aβ Free Nerve Ending sensory neuron in glabrous skin that will be described elsewhere (*Ptgfr^CreER^*) (Luo et al., 2009; Bai et al., 2015; Zheng et al., 2019)(Figure 3B; see Methods).

**Figure 3.**
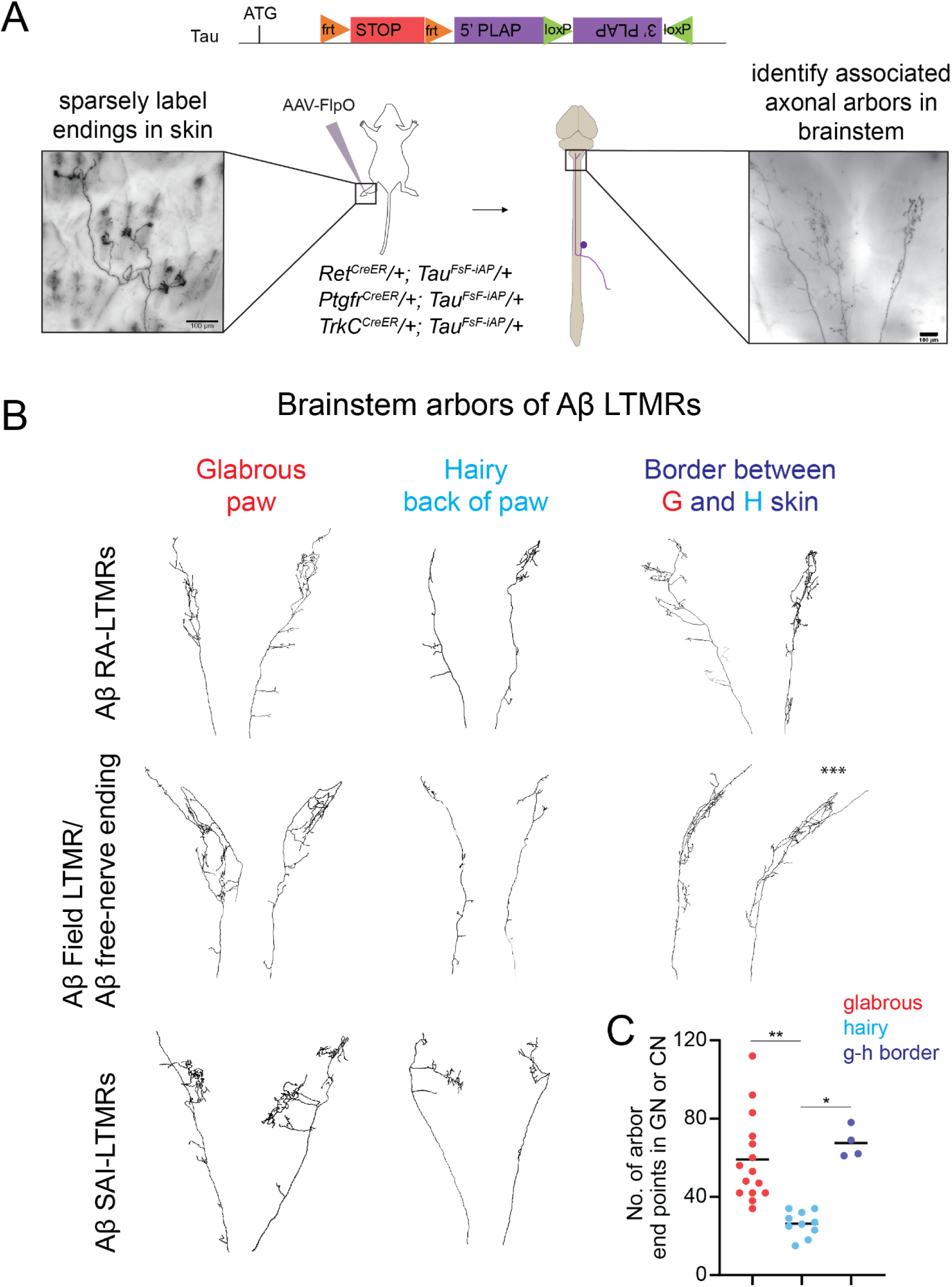
Target dependence of Aβ-LTMR central arbor morphologies. **A**. Sparse neuron labeling using alkaline phosphatase (AP) reporters allows for examination of the peripheral and brainstem arbor morphologies of individual neurons. Generation of an AP reporter mouse in the *Tau (Mapt)* locus allows intersectional labeling of Aβ-LTMR types with specific CreER lines and AAV2/1-hSyn-FlpO virus injection to hairy or glabrous skin. Animals with sparse labeling are analyzed to relate peripheral arbor morphologies (left) to central arbor anatomies in the gracile or cuneate nuclei (right). **B**. Representative examples of reconstructed brainstem arbors of Aβ RAI-LTMRs (top), Aβ field/free-LTMRs (middle), or Aβ SAI-LTMRs (bottom) that innervate glabrous or hairy paw skin. Across genetic and functional classes, glabrous skin innervating neurons consistently form more elaborate axonal arbors in the gracile or cuneate nuclei than most hairy skin innervating neurons. Neurons innervating hair follicles that lie close to the border with glabrous skin (<0.5 mm) or that span both types of skin form complex arbors that are indistinguishable from those formed by glabrous skin innervating neurons (right). The neuron whose peripheral arbor spanned both glabrous and hairy skin is indicated with asterisks (***); the other three neurons innervated hair follicles close to glabrous skin. **C**. Quantification of the total number of end points formed by the central arbors of Aβ-LTMRs in the brainstem. ** indicates p < 0.0005 and * indicates p < 0.01, by Welch’s t-test.

We reconstructed the entire axonal arbors of each Aβ-LTMR type in the GN or CN for adult neurons that innervate paw glabrous skin, paw hairy skin, or trunk hairy skin (Figure 3 and Supplemental Figure 3). For each Aβ-LTMR reconstruction, we quantified the number of arbor end points and the total length of axon collaterals in the GN and CN volumes. We found that paw glabrous skin-innervating neurons form elaborate central arbors in the GN and CN that dwarf those formed by trunk level LTMRs and by the majority of paw hairy skin-innervating LTMRs (Figure 3B-C, Supplemental Figure 3). Intriguingly, the central arbors of a small number of paw hairy skin-innervating LTMRs were equally as complex as those of glabrous skin-innervating LTMRs (Figure 3B-C). To determine if this variability in hairy LTMR central arbor structure is explained by LTMR subtype or location, we identified the exact position and morphology of each neuron’s peripheral arbor within the hairy skin of the paw. We detected no relationship between the genetic class of neurons and the size of their central arbors, or between the surface area of a neuron’s peripheral arbor and the complexity of its central arbor (Supplemental Figure 3). There was also no detectable difference when comparing the overall structure of neurons targeting the GN or CN. Instead, we found that hairy skin-innervating neurons with peripheral arbors located at the border between paw glabrous skin and paw hairy skin (<0.5 mm from the edge of the paw after dissection) form arbors in the GN or CN that are significantly more elaborate than those formed by LTMRs innervating hairy skin >0.5 mm from the edge of the paw (Figure 3B-C, Supplemental Figure 3). Interestingly, one of these “edge of paw” neurons formed peripheral endings that straddle both hairy and glabrous skin, suggesting that the innervation territories of some Aβ-LTMRs are not restricted to one type of skin. This neuron formed a complex central arbor in the brainstem that resembled those of pure glabrous neurons and the other three edge of paw neurons (Figure 3C, arbor indicated by asterisks). Together, this demonstrates that the proximity of an Aβ-LTMR’s peripheral arbor to glabrous skin predicts the complexity of its axonal outputs to brainstem across multiple genetically and morphologically distinct LTMR subclasses.

We next investigated how Aβ-LTMR central arbors change over development. Low dose tamoxifen treatment of *Ret^CreER^; Brn3a^cKOAP^* mice sparsely labels Aβ RA-LTMRs throughout the animal, the majority of which innervate trunk hairy skin. We examined how the central arbor structure of this Aβ LTMR subtype changes between early postnatal and adulthood timepoints. While this sparse labeling strategy was not effective for visualizing the central arbors of paw glabrous skin-innervating neurons, we found that P5-P10 hairy skin-innervating and trunk-level Aβ RA-LTMRs form significantly more complex axonal arbors in the GN or CN compared to their adult counterparts when quantifying the total number of end points, axon collateral lengths, and the number of axon collaterals (Supplemental Figure 3 and data not shown). This demonstrates that refinement of the central projections of Aβ RA-LTMRs occurs over postnatal development.

The dramatic structural differences observed for the central arbors of Aβ-LTMRs innervating different skin regions and at different ages suggested that the number of synapses of individual Aβ-LTMRs in the GN or CN may also reflect the type of skin they innervate and change as a function of developmental age. Therefore, we next explored the relationship between Aβ-LTMR skin innervation sites and their histologically-defined synapses in the brainstem. Presynaptic terminals of Aβ-LTMRs were labeled with Synaptophysin-tdTomato or Synaptophysin-mScarlet fusion proteins, hereafter referred to as syn-fluorescent protein (syn-FP), by injecting AAVs into the glabrous or hairy skin of the paw (Figure 4A). We quantified the number of syn-FP^+^ puncta in the GN and CN and the number of labeled LTMR axons entering the brainstem to calculate average synapse numbers per LTMR. In GN and CN sections co-stained with synaptic markers, the majority of syn-FP^+^ puncta were found in apposition to Homer1, a post-synaptic marker for excitatory synapses, and co-localized with vGlut1, a presynaptic marker for Aβ-LTMRs (Figure 4A and Supplemental Figure 4). This analysis revealed that individual adult forepaw glabrous skin innervating neurons form significantly more synapses in the CN than adult forepaw hairy skin innervating neurons (Figure 4B p<0.05 by Welch’s t-test, 176.4 synapses/neuron for glabrous forepaw Aβ-LTMRs, 96.2 synapses/neuron for hairy forepaw Aβ-LTMRs). Similarly, adult hindpaw glabrous skin innervating neurons form 2.5-fold more synapses in the GN than adult hindpaw hairy skin innervating neurons (Figure 4C, p<0.01 by Welch’s t-test, 79.5 synapses/neuron for glabrous hindpaw Aβ-LTMRs, 31.8 synapses/neuron for hairy hindpaw Aβ-LTMRs). Pre-synaptic boutons were found both at arbor end points and along the length of axonal fibers, at an equivalent rate in glabrous and hairy neurons (Supplemental Figure 4).

**Figure 4.**
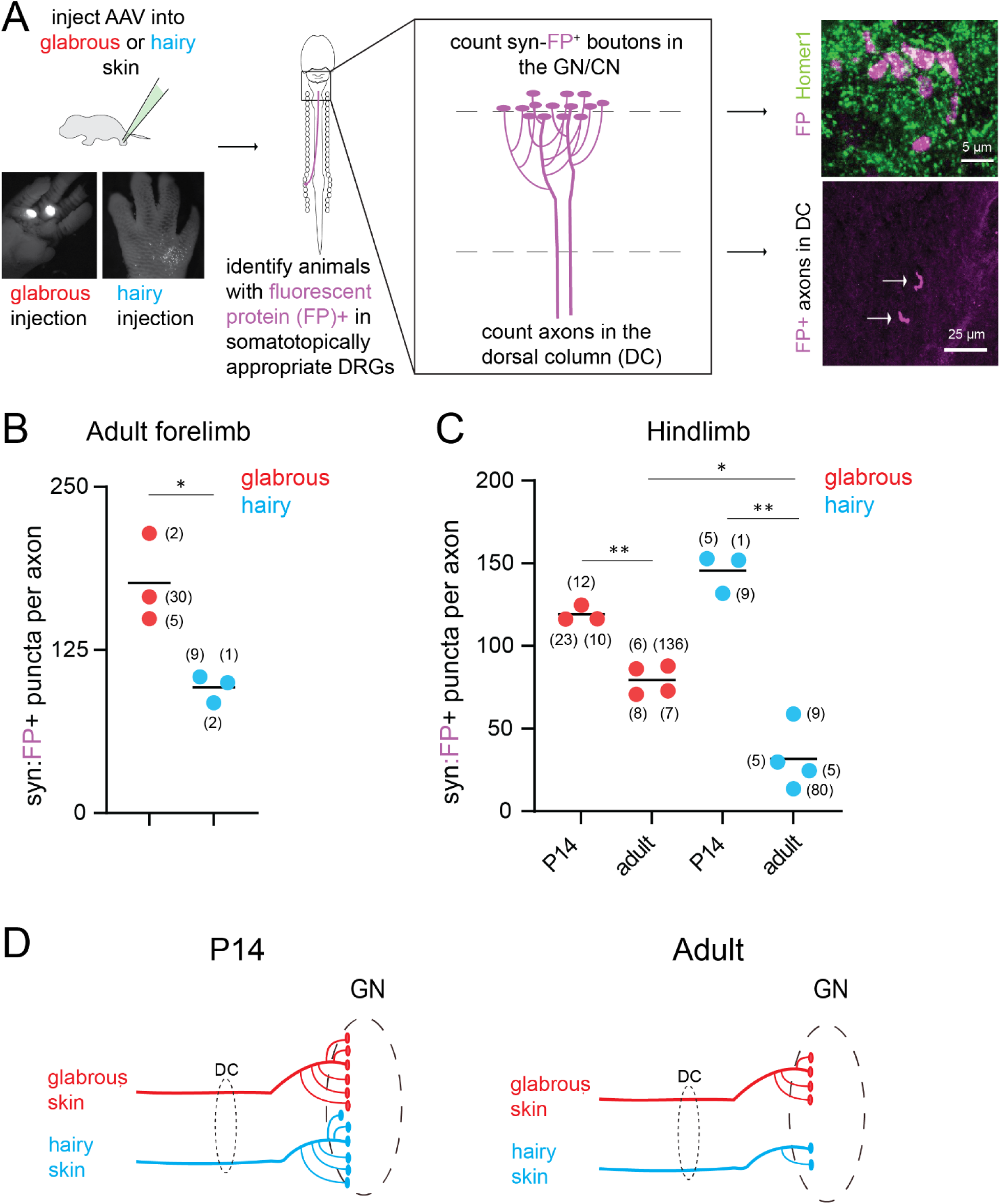
Relationship between skin target identity and Aβ-LTMR brainstem synapses over development. **A**. (left) AAV virus injection to glabrous or hairy paw skin allows for skin type-specific labeling of Aβ-LTMR central projections with synaptophysin-tdTomato or synaptophysin-mScarlet, hereafter referred to as synaptophysin-FP (fluorescent protein). Animals were screened following tissue collection to confirm that viral transduction was restricted to DRG at somatotopically appropriate segmental levels (C7-C8 for forepaw injections, L3-L5 for hindpaw injections). The gracile and cuneate nuclei were sectioned and stained for fluorescent proteins and synaptic markers. (right) Representative example of labeled boutons apposed to the excitatory postsynaptic marker Homer1 in the gracile nucleus (top). Both hairy and glabrous skin-innervating Aβ-LTMRs form syn-FP^+^ boutons apposed to Homer1^+^ puncta in the gracile or cuneate nuclei. Representative example of labeled axons (arrows) ascending the dorsal column, in a transverse section of the cervical spinal cord (bottom). Arrows point to individual axons. **B-C**. Quantification of the average number of FP+ pre-synaptic boutons formed by Aβ-LTMRs innervating different skin targets. Each data point represents one animal; the number of axons identified in each animal is shown in parentheses. **B**. Adult forepaw glabrous skin-innervating Aβ-LTMRs form more synapses than forepaw hairy skin-innervating Aβ-LTMRs in the cuneate nuclei (*p<0.05, Welch’s t-test). **C**. Hind paw glabrous skin-innervating Aβ-LTMRs and hindlimb hairy skin-innervating Aβ-LTMRs form more pre-synaptic boutons in the gracile nuclei at P14 than in adulthood (**p<0.001, Welch’s t-test). However, developmental synapse loss is more pronounced in hairy skin-innervating Aβ-LTMRs, such that at adult time points, paw glabrous skin Aβ-LTMRs form significantly more pre-synaptic boutons in the gracile nuclei than paw hairy skin Aβ-LTMRs (*p<0.01, Welch’s t-test). **D**. Cartoon illustrating a model for skin type-dependent synaptic refinement in Aβ-LTMRs. At early postnatal ages, Aβ-LTMRs that innervate hairy and glabrous skin form comparable numbers of pre-synaptic specializations in the brainstem. Between P14 and P50, Aβ-LTMRs undergo synaptic bouton loss in a skin type-dependent manner, such that in adults, Aβ-LTMRs innervating glabrous skin form more pre-synaptic boutons in the brainstem than those that innervate hairy skin.

To determine when these synaptic differences in Aβ-LTMR central arbors emerge over development, we focused on the hindlimb and injected AAV-syn-FP into hindpaw glabrous skin, hindpaw hairy skin, or thigh hairy skin to label Aβ-LTMRs in young animals. Importantly, at P14, both hindlimb hairy skin- and glabrous skin-innervating Aβ-LTMRs form more synapses in the GN than observed in adulthood (Figure 4B). However, this difference is more pronounced for hindpaw hairy skin-innervating Aβ-LTMRs, which exhibited a ~75% loss of synapses per neuron between P14 and adulthood. Indeed, at P14, hindlimb hairy skin-innervating Aβ-LTMRs form slightly more synapses in the GN than their glabrous counterparts, in contrast to adult timepoints where glabrous paw Aβ-LTMRs form ~2.5-fold more synapses than hairy paw Aβ-LTMRs (Figure 4B).

To further explore the functional consequences of maturation of LTMR inputs to the brainstem over development, we used the GN functional imaging platform to analyze receptive fields of individual touch responsive GN thalamic projection neurons at postnatal and adult timepoints. We found that receptive fields decline in size over time, consistent with developmental refinement and possible pruning of synaptic connections. This is apparent in the percentage of the population of thalamic projection neurons with multiregional receptive fields, defined as GN thalamic projection neurons that responded to touch on two or more of the following body regions: glabrous paw, hairy paw, hairy thigh, hairy back (Supplemental Figure 4). Fewer GN thalamic projection neurons with multiregional receptive fields were observed as development progressed (Supplemental Figure 4). Glabrous expansion could occur through an increase in the number of neurons responding to glabrous touch, or a depletion of those responding to hairy skin stimulation. We observed a decline in the percentage of the GN thalamic projection neuron population that responded to hairy skin, but not glabrous skin (Supplemental Figure 4).

Taken together, these histological and functional analyses reveal that paw glabrous skin-innervating Aβ-LTMRs exhibit more elaborate central morphologies and more synapses in the brainstem compared to paw hairy skin-innervating LTMRs and that these differences emerge over postnatal development (Figure 4D). Moreover, these morphological and synaptic differences occur in conjunction with developmental changes in GN thalamic projection neuron receptive field properties. These findings thus support a model in which developmental refinement of LTMR synaptic connectivity within the brainstem produces disproportionate emphasis on glabrous skin surfaces in the GN and CN thalamic projection neuron populations.

### Optogenetic receptive field mapping shows that single LTMR stimulation in glabrous skin more powerfully excites GN neurons

The functional connectivity of neurons in the visual system has been defined at single photoreceptor resolution through approaches that relate the history of spiking to a white noise stimulus in a neuron’s receptive field (Field et al., 2010). We applied this approach to our analysis of the somatosensory system by expressing the light-gated ion channel ReaChR (Hooks et al., 2015) in peripheral neurons and explored the synaptic mechanisms of disproportionate body representation in the brainstem. Whereas mechanical stimuli that reliably drive spiking inevitably excite many Aβ LTMRs in skin, optogenetic approaches have the potential to selectively and reliably excite single LTMRs. We therefore developed methods for making electrophysiological recordings of touch responsive neurons in the DRG and brainstem while delivering precise, focal optogenetic stimulation to peripheral neuron terminals in skin.

We constructed an optomechanical device capable of rapidly and precisely exciting the endings of neurons in the skin of *Cdx2-Cre; Avil^FlpO^*; *Rosa26^LsL-FsF-ReaChR^* mice, which express ReaChR in all sensory neurons below cervical level C2 (Figure 5A). We recorded from L4 DRG in adult animals with an MEA and both mechanically and optically stimulated the ipsilateral thigh and paw. In the representative example DRG unit shown in Figure 5B, we first mapped the mechanical receptive field to a digit on the paw that exhibited a rapidly adapting response to a force-controlled indentation (Figure 5B, top right). We next optically stimulated ReaChR expressing sensory neuron terminals in the skin with focused pulses (300 microseconds) of laser light, moving the stimulus to a new location randomly chosen from across the entire paw region, every 1.1 msec (Figure 5B, bottom). We then used reverse correlation analysis to recover a time-varying spatial receptive field for this unit through computation of the spike-triggered average (Figure 5B, bottom). The optical receptive fields generated in this manner resemble the mechanosensory terminal fields of individual LTMRs and overlapped with their mechanical receptive fields (n=11 neurons with targeted mechanical stimulation, data not shown). Importantly, Aβ-LTMRs in both hairy and glabrous skin were excited by optical stimulation with equal probabilities and latencies (Figure 5C, data not shown), making it possible to directly compare receptive fields on glabrous and hairy skin.

**Figure 5.**
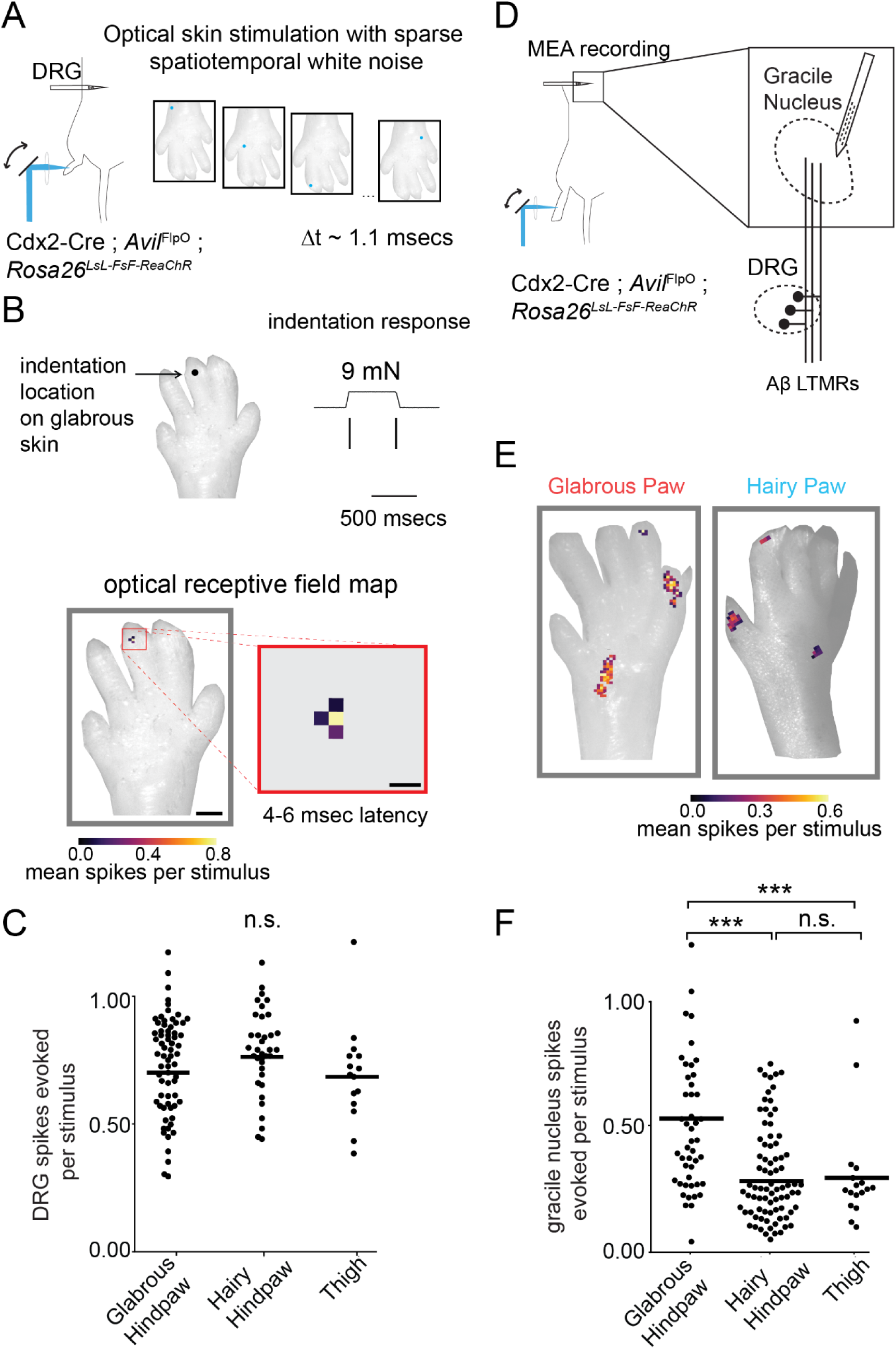
Aβ LTMRs that innervate glabrous skin more powerfully excite central tactile neurons in the gracile nucleus than Aβ LTMRs that innervate hairy skin. **A**. Simultaneous *in vivo* recording from tens of LTMRs with cell bodies in L4 DRG was paired with optical stimulation of skin with sparse spatiotemporal white noise. Pulses of laser light (300 microseconds, 20 mW) were directed by a pair of galvo mirrors and focused through an f-theta lens onto the skin. Laser pulses were delivered every 1.1 msec for a total of 1.3 million pulses per session. Experimental mice expressed ReaChR in all cutaneous sensory neurons. **B**. Representative single LTMR unit recorded *in vivo* with a MEA in the L4 DRG. (top) Spiking response to mechanical indentation for an Aβ RA-LTMR. (bottom) Representative spike-triggered average responses describing the spatiotemporal receptive field of the same RA-LTMR computed through reverse correlation. Spatial receptive fields produced from the spike-triggered average by integrating over the response window (4-6 msec) are overlayed over an image of the paw, and only values above the noise threshold are displayed. LTMRs have spatiotemporally simple receptive fields in skin that align with mechanical receptive fields and have dimensions similar to terminal endings. Scale bars are 1 mm and 200 μm. **C**. The mean number of spikes evoked by a stimulus to the center of a LTMRs optogenetic receptive field, computed by integrating the conditional probability of a spike over the 3 −6 msec following a light pulse. An equivalent number of LTMR spikes were evoked by stimulating optogenetic RFs in glabrous hindpaw skin, hairy hindpaw skin, and thigh skin (following hair removal). **D**. Strategy for exploring the functional connectivity of the gracile nucleus at single LTMR resolution. Optogenetic receptive fields of touch sensitive neurons in the gracile nucleus were characterized by computing the spike-triggered average of a sparse spatiotemporal white noise stimulus presented to the skin of mice that expressed ReaChR in all sensory neurons. **E**. Representative units recorded from the gracile nucleus with receptive fields on the glabrous (left) or hairy hindpaw skin (right). Gracile units are complex yet composed of spatially and temporally separable “hotspots” in their RFs, indicating convergence of multiple LTMRs onto central touch neurons in the gracile nucleus. **F**. The number of spikes evoked in a gracile nucleus neuron by a single optical pulse to “hotspots” comprising their RFs. Spots reminiscent of single LTMR RFs are present for all skin targets, and optical stimulation of glabrous skin innervating sensory neurons is more effective at driving spikes in postsynaptic neurons in brainstem than hairy skin optical stimulation (one-way ANOVA, p < 10^-13^, *** indicates Bonferroni corrected p < 10^-6^ via Mann Whitney U Test).

Next, a sparse white noise optical stimulus was used to excite either hindpaw glabrous skin-innervating neurons, hindpaw hairy skin-innervating neurons, or thigh-innervating neurons while recording from the GN of the brainstem using an MEA targeted to the hindpaw region (Figure 5D). We found that the majority of GN units were both mechanically and optically responsive. In contrast to the LTMRs recorded in the DRG (Figure 5A-C), however, GN unit spike-triggered averages demonstrated complex spatiotemporal receptive fields with multiple, discontinuous RF “spots” that were reminiscent of individual LTMR spatial RFs and staggered with respect to one another in time (Figure 5E, Supplemental Figure 5C-D). RF spots had similar strength and spatial receptive fields in control experiments where a fraction of Aβ-LTMRs expressed ReaChR, though the number of spatiotemporally isolated spots identified declined, as expected if each spot reflects the contribution of a single LTMR input to a GN neuron’s RF (Supplemental Figure 5E, F). Furthermore, the recorded latencies were consistent with direct synaptic input from Aβ-LTMRs after accounting for the time required to transit the dorsal column, the delay across the synapse, and the time for spike generation, but may also include indirect input from the post-synaptic dorsal column pathway.

Examination of the time-projected spatial receptive fields of optically-evoked GN units demonstrates that stimulation of a single “RF spot” – which likely corresponds to a single Aβ-LTMR input – in glabrous skin is more effective at driving spiking in GN neurons than a similar stimulus applied to a hairy skin spatial subunit (Figure 5E, F). This effect was consistent across 120 spatiotemporally well-isolated RF spots identified across eight animals (Figure 5F). Because the optical stimuli used for these GN maps are equally effective at exciting hairy and glabrous skin LTMRs, based on our DRG recordings (Figure 5C), this finding demonstrates that stimulation of individual glabrous skin-innervating Aβ-LTMRs more strongly evokes spiking in GN neurons. These findings suggest a model in which differential LTMR synaptic drive underlies the expansion of glabrous skin representation in the GN.

### Fiber fraction measurements of synaptic strength and convergence reveal skin region dependent differences in LTMR inputs to thalamic PNs in the GN

Our histological synaptic analysis (Figure 4) and optogenetic receptive field measurements (Figure 5) suggest that glabrous skin innervating LTMRs make more numerous and more powerful synaptic connections onto GN neurons compared to hairy skin innervating LTMRs. However, the *in vivo* optogenetic RF experiments provide only an indirect measure of synapse strength that lacks cellular specificity. To directly characterize synaptic connections between glabrous skin Aβ-LTMRs or hairy skin Aβ-LTMRs and GN thalamic projection neurons, we made whole-cell patch clamp recordings from GN thalamic projection neurons in acute brainstem slices. The relative strength of glabrous skin and hairy skin innervating Aβ-LTMR synapses were measured by optogenetically exciting single presynaptic axons in animals that expressed the ReaChR in RA-LTMRs that innervate glabrous or hairy skin, respectively. To measure the total ascending synaptic input and produce corresponding fiber fraction estimates of the convergence of presynaptic inputs onto GN thalamic projection neurons, we also electrically stimulated the dorsal column with a bipolar electrode (Hooks and Chen, 2006).

Glabrous skin Aβ RA-LTMRs that innervate Meissner corpuscles and hairy skin Aδ-LTMRs are both labeled in *Ntrk2^CreER^; Avil^FlpO^; R26^LSL-FsF-ReaChR::mCitrine^* mice. However, hairy skin Aδ-LTMRs do not project up the dorsal column, and thus their axons are absent in horizontal acute brain slices that include the GN and dorsal column. Therefore, we used this genetic labeling strategy to preferentially activate glabrous skin Aβ RA-LTMR inputs to the GN using optogenetics (Supplemental Figure 6). To achieve corresponding selective genetic access to hairy skin Aβ RA-LTMR inputs in brainstem slices, we noted that Pacinian corpuscle innervating Aβ RA2-LTMRs, Meissner corpuscle innervating Aβ-LTMRs, and hairy skin Aβ RA-LTMRs are all labeled by tamoxifen administration at Ell.5 in *Ret^CreER^* mice, but hairy skin Aβ RA-LTMRs are the only population within this group that are not labeled by *Pvalb^FlpO^* (Supplemental Figure 6). Serendipitously, the FRT site within the *neo* selection cassette flanking ChR2 in Cre-dependent ChR2 transgenic mice *R26^LsL-FRT-ChR2::YFP-attB-FRT-Neo-attP^* (Ai32D) remains unexcised, allowing for a Cre ON, Flp OFF subtractive strategy whereby Aβ RA-LTMRs that form lanceolate endings in hairy skin were selectively labeled in *Ret^CreER^; Pvalb^FlpO^; R6^LsL-F-ChR2::YFP-F^* mice (Supplemental Figure 6). Therefore, we used *Ntrk2^CreER^; Avil^FlpO^; R26^LsL-FsF-ReaChR::mCitrine^* mice and *Ref^reER^; Pvalb^FlpO^; R26^LsL-F-ChR2::YFP-F^* mice to selectively express an opsin in the GN terminals of either glabrous skin- or hairy skin-innervating Aβ RA-LTMRs, respectively (Figure 6A; Supplemental Figure 6).

**Figure 6.**
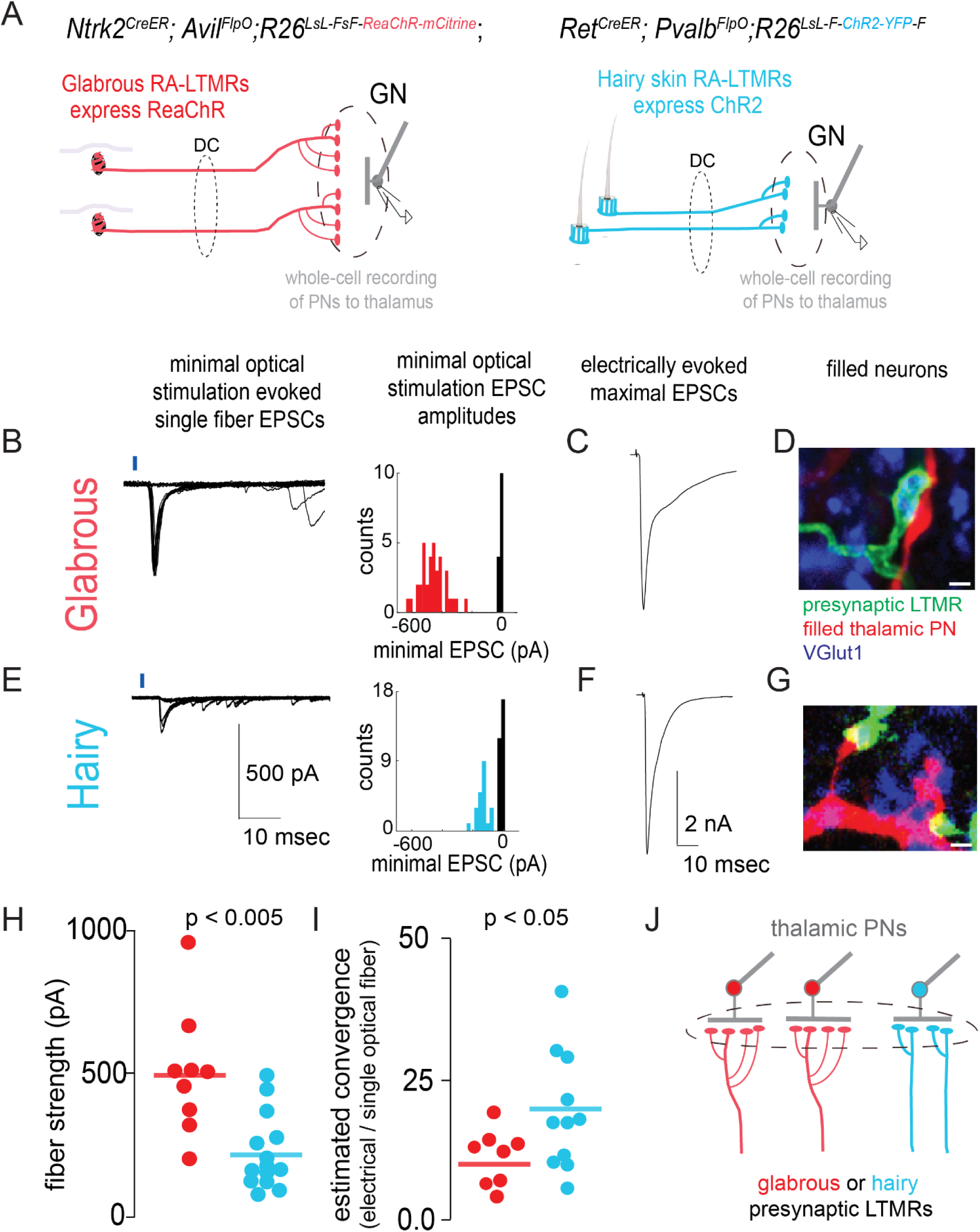
Synaptic strength and functional convergence of glabrous skin- and hairy skin-innervating LTMR synapses onto thalamic projection neurons in the brainstem. **A.** Genetic strategies for assessing synaptic connections between glabrous skin- or hairy skin-innervating LTMRs and thalamic projection neurons in acute brainstem slices. (left) Glabrous skin Aβ RA-LTMRs that send ascending projections through the dorsal column and synapse in the gracile nucleus express ReaChR in *Ntrk2^CreER^; Avil^FlpO^; R26^LsL-FsF-ReaChR::mCitrine^* mice. The vast majority of hairy skin-innervating neurons labeled under this strategy do not ascend the dorsal column and are not captured in acute brainstem slices (see Supplemental Figure 5). (right) Both glabrous skin- and hairy skin-innervating Aβ RA-LTMRs are labeled by *Ret^CreER^* with embryonic tamoxifen at E10.5-E11.5, but hairy skin Aβ RA-LTMRs do not express *Pvalb^FlpO^*. Thus, hairy skin-innervating Aβ RA-LTMRs exclusively express ChR2 using a Cre ON, Flp OFF strategy. **B**. Representative whole-cell recording from a retrogradely labeled thalamic projection neuron in the gracile nucleus. (left) Minimal optical stimulation of glabrous Aβ RA-LTMRs axons in the dorsal column (1-4 mm from the gracile nucleus) isolated putative single fiber-evoked EPSCs as stereotyped currents accompanied by failures. (right) Histograms of EPSC amplitudes for these representative recordings are consistent with single fiber stimulation. **C**. EPSC evoked by maximal stimulation of axons ascending the dorsal column, from the same recording as B. **D**. The biocytin fill of a recorded cell (red) shows apposition to VGlut1^+^ (blue), ReaChR::mCitrine+ (green) presynaptic boutons, consistent with a direct synaptic connection (scale bar: 1 μm). **E**. Representative whole-cell recording from a retrogradely labeled thalamic projection neuron in the gracile nucleus postsynaptic to hairy skin innervating Aβ RA-LTMRs. Minimal optical stimulation isolates a single short-latency input (left) with trial-to-trial amplitudes consistent with successes and failures of a single fiber. **F**. EPSC evoked by maximal stimulation of all axons ascending the dorsal column in the same recording as D. **G**. Biocytin filled dendrites (red) are apposed to VGlut1+ (blue), ReaChR::mCitrine+ (green) presynaptic boutons, consistent with a direct synaptic connection (scale bar: 1 μm). **H**. Optogenetic activation of individual glabrous skin innervating Aβ RA-LTMRs neurons evokes a larger EPSC than activation of individual hairy skin-innervating Aβ RA-LTMRs (p < .005, Mann Whitney U test). **I**. The estimated functional convergence for thalamic projection neurons receiving synaptic input from glabrous skin- (red) and hairy skin-innervating (blue) Aβ RA-LTMRs computed by dividing the maximal EPSC evoked by electrical stimulation of the dorsal column by the genotype-specific single fiber strength. **J**. A model of differential synaptic expansion consistent with synaptic strength and convergence measurements presented in (A-I). Glabrous skin-innervating LTMRs have more synaptic release sites per neuron, and fewer presynaptic fibers converge onto postsynaptic thalamic projection neurons than LTMRs that innervate hairy skin. Glabrous LTMR circuits thus have a lower functional convergence ratio while maintaining comparable synaptic input to thalamic PNs, resulting in synaptic expansion that enlarges glabrous skin representation.

GN thalamic projection neurons in horizontal brainstem slices of *Ntrk2^CreER^; Avil^FlpO^; R26^LsL-F-ChR2::YFP-F^* mice and *Ref^CreER^; Pvalb^FlpO^; R26^LsL-F-ChR2::YFP-F^* mice were retrogradely labeled from the thalamus with CTB-647 and visually targeted for whole cell patch clamp recordings.

Using this preparation, saturating light pulses (150 mW/mm^2^, 1-2 msec in duration) applied to axons in the dorsal column produced short latency EPSCs in a fraction of GN thalamic projection neurons held at −70 mV, which is near the inhibitory reversal potential, establishing a synaptic connection between glabrous skin or hairy skin Aβ RA-LTMRs and the recorded GN thalamic projection neuron. Next, the area of illumination was restricted, the light power was reduced, the light pulse shortened, and the illuminated region shifted caudally to optogenetically activate individual LTMR axons (Litvina and Chen, 2017). Under these conditions, minimal optical stimulation of both glabrous skin-innervating Aβ RA-LTMRs and hairy skin-innervating Aβ RA-LTMRs produced short-latency unitary synaptic EPSCs interspersed with failures (Figure 6B, E, left panels). EPSC waveforms were stereotyped, and histograms of the evoked synaptic current amplitudes across multiple trials were bimodal, consistent with single fiber isolation (Figure 6B, E, right panels). Maximal EPSCs evoked through saturating electrical stimulation of dorsal column axons were of comparable amplitude across recordings from GN thalamic projection neurons postsynaptic to glabrous skin- and hairy skin-innervating LTMRs (Figure 6C, F, Mann Whitney U test p > 0.05, data not shown). Immunohistochemical visualization of genetically labeled LTMRs and filled, recorded GN thalamic projection neurons revealed regions of presynaptic (fluorescent protein and VGlut1+) and postsynaptic, dendritic apposition in all cases where synaptic connections were present, consistent with a monosynaptic connection (Figure 6D,G).

These GN whole-cell recordings revealed that glabrous skin-innervating Aβ RA-LTMR single fiber strengths are approximately two-fold larger than hairy Aβ RA-LTMR fiber strengths (Figure 6H; Mann Whitney U test, p < 0.005). Fiber fractions were computed for GN thalamic projection neurons postsynaptic to glabrous skin- or hairy skin-innervating Aβ RA-LTMRs by dividing the minimal optogenetic single fiber EPSC by the maximal electrically-evoked EPSC (Litvina and Chen, 2017). This reciprocal of the heterotypic fiber fraction is an estimate of the total number of presynaptic ascending neurons that converge onto individual GN thalamic projection neurons. Importantly, our electrical stimulation findings show that “glabrous skin” and “hairy skin” GN thalamic projection neurons receive comparable total synaptic input.

These findings support a model where thalamic projection neurons receive similar amounts of total synaptic input, but glabrous skin-innervating LTMRs capture a disproportionate share of the projection neuron population by forming more presynaptic inputs with stronger single-fiber strengths, and displaying less convergence relative to hairy skin-innervating LTMRs (Figure 6J).

### Disproportionate glabrous skin representation is maintained in mice that do not sense gentle touch

In the visual system, both spontaneous and visually evoked patterns of retinal activity contribute to pruning of retinal ganglion cell synapses in the thalamus during postnatal development (Hooks and Chen 2006). Therefore, we next considered the possibility that touch-evoked LTMR activity might be an important force influencing synapse formation in the GN and CN, and that differences in tactile experiences might shape the proportionate representation of the body. To determine if mechanically evoked activity in LTMRs shapes the relative strength of Aβ-LTMR synapses in the brainstem, we analyzed conditional knock-out animals in which Piezo2, the principal mechano-sensitive ion channel for LTMRs (Ranade et al., 2014), is deleted using the *Cdx2-Cre* line, which drives recombination in all DRG neurons below cervical level C2 as well as in glia and epidermal cells of the hindpaw. *Cdx2-Cre; Piezo2^flox/flox^* and *Cdx2-Cre; Piezo2^flox/nuil^* animals survive to adulthood, but display severe proprioceptive defects, similar to those reported for *Hoxb8-Cre; Piezo2* cKO animals (Woo et al., 2015; Murthy et al., 2018). *In situ* hybridization experiments in *Cdx2-Cre; Piezo2^flox/flox^* animals demonstrate efficient deletion of the floxed coding sequence in lumbar-level DRG neurons, with no detectable difference in DRG size or neuron numbers (Supplemental Figure 7). All expected mechanosensory neuron peripheral ending types are present in glabrous and hairy skin of *Cdx2-Cre; Piezo2^flox/flox^* animals (data not shown). We used these mice to assess touch sensitivity and optical excitability of Aβ LTMRs that lack *Piezo2*, with the goal of comparing optical receptive fields from different skin regions in GN neurons in brainstem.

In wild type animals that express ReaChR in all sensory neurons, the majority of DRG units that spike in response to optical stimulation of skin also respond to mechanical brushing of the skin (Figure 7A). In contrast, touch evoked responses are nearly abolished in *Cdx2-Cre; Piezo2^flox/flox^; Avil^FlpO^; R26^-FsF-ReaChR^* animals, while optical responses to skin stimulation persisted (Figure 7B). Importantly, the number of spikes evoked by optical stimulation within DRG sensory neurons’ RFs were equivalent between *Piezo2* conditional mutants and control animals (Figure 7C, p = 0.49 Mann Whitney U test).

**Figure 7.**
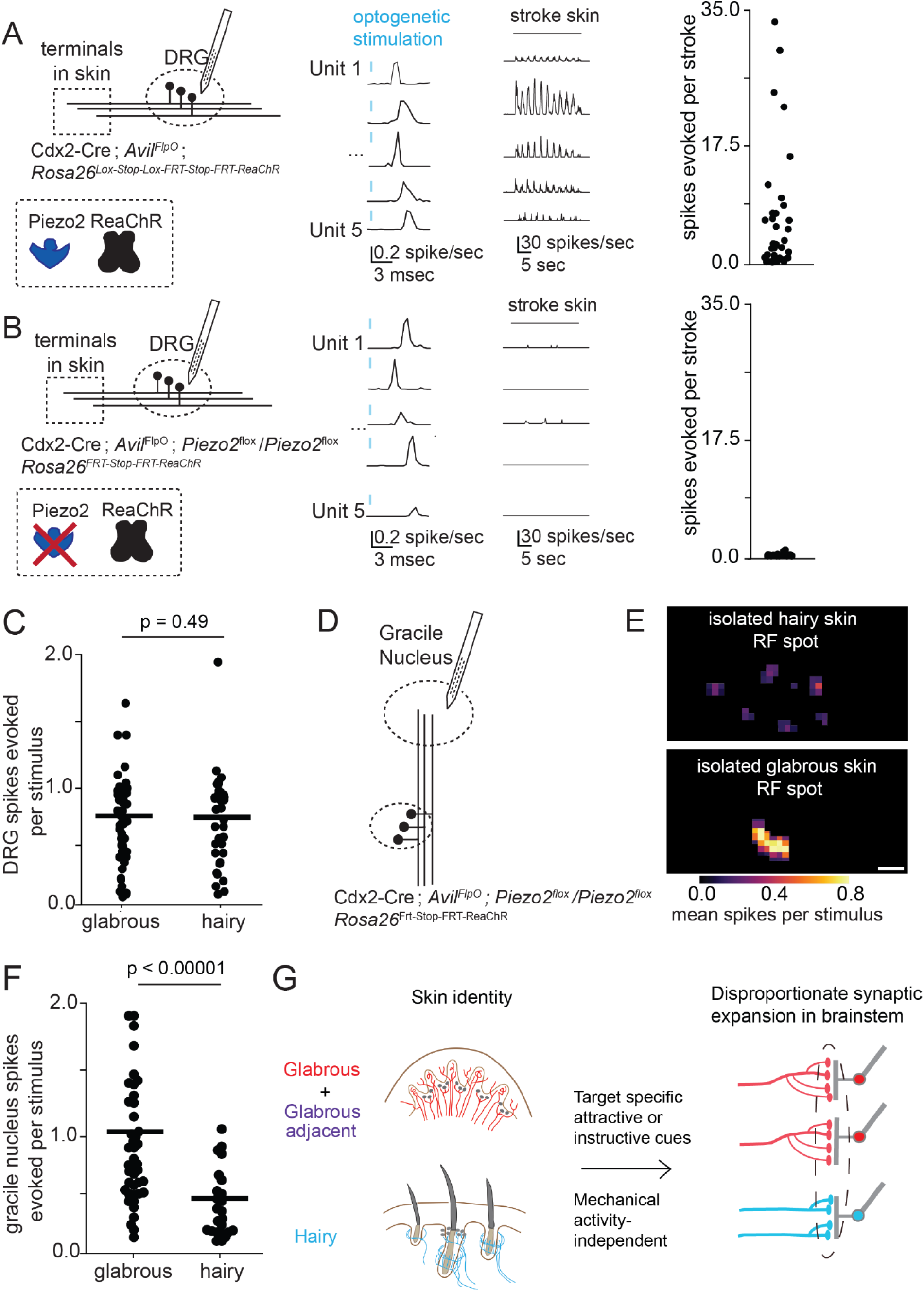
Disproportionate glabrous skin representation is maintained in mice that do not sense gentle touch. **A**. (left) MEA recordings from L4 DRGs in wild-type mice expressing ReaChR in all sensory neurons. Units that responded to optical stimulation of the skin with short-latency spikes (center) also showed robust responses to stroke on the skin (right). **B**. MEA recordings from L4 DRG in *Piezo2* conditional knockout (P2 cKO) animals that express ReaChR in all sensory neurons (*Cdx2-Cre; Piezo2^flox/flox^; Avil^FlpO^; R26^FsF-ReaChR^*). Optical stimulation of the skin produces short-latency spikes (center), as seen in control mice (A), however few to no responses to mechanical stimuli were observed in Piezo2 mutants (right). **C**. Optical stimulation of the receptive field of Piezo2 cKO DRG neurons was equally effective for neurons innervating glabrous and hairy skin (p=0.49, Mann-Whitney U test). **D**. Strategy for exploring the functional connectivity of the gracile nucleus at single LTMR resolution in Piezo2 cKO animals. Optogenetic receptive fields of touch sensitive neurons in the gracile nucleus were characterized by computing the spike-triggered average of a sparse spatiotemporal white noise stimulus presented to the skin of mice that expressed ReaChR in all sensory neurons. **E**. Representative well-isolated receptive field subunits of gracile neurons computed after hairy skin stimulation (top) or glabrous skin stimulation (bottom). These subunits are consistent with the anatomy of LTMR terminals in the skin of wild type animals. **F**. The number spikes evoked in units recorded from the gracile nucleus following optical stimulation of receptive field subunits in glabrous hindpaw skin, hairy hindpaw skin, and thigh skin (after hair removal). Spots reminiscent of single LTMR ending anatomies are present for all skin targets, and all units analyzed were mechanically unresponsive. **G**. Disproportionate expansion of glabrous skin representation in the brain is driven by the skin region innervated by sensory neurons. As tactile-evoked responses are not required for the developmental expansion of glabrous skin representation, we propose that the ventral paw/glabrous skin either i) provides an instructive signal to developing neurons to form more synaptic connections in the brainstem or ii) generates cues that attract neurons that are preprogrammed to make more synaptic connections in the brainstem.

Therefore, we made extracellular MEA recordings from the GN in *Cdx2-Cre; Piezo2^flox/flox^; Avif^lpO^; R26^FsF-ReaChR^* mutants (similar to the experiments on *Cdx2-Cre; Avil^FlpO^; R26^LsL-FsF-ReaChR^* mice presented in Figure 3) while presenting a sparse white noise optical stimulus to the skin (as depicted in Figure 5D). Spatiotemporally well-isolated single RFs with dimensions reminiscent of individual LTMRs were recovered from spike-triggered averages. These isolated RFs represent convergent, and most likely single LTMR inputs to GN neurons (Figure 7E). Similar to control animals, we found that stimulation of RFs on glabrous skin more strongly excited brainstem neurons than comparable stimuli applied to hairy skin in the *Piezo2* mutant mice that lack mechanically-evoked LTMR activity (Figure 7E, F). Together, these findings indicate that the disproportionate strength of glabrous skin responses in the brainstem can be established independent of mechanically-evoked activity.

## Discussion

Mammalian nervous systems are embedded in a fascinating diversity of body forms that in turn shape their structure, as evidenced by the fact that nearly half of the mouse brain contains neurons that are topographically organized with respect to the body (Wang et al., 2020). Glabrous skin responsive neurons occupy disproportionately large regions of somatosensory cortex in primates, cats, squirrels, and bats, suggesting a common mechanism might unite these observations (Sur et al., 1978; Sur et al., 1980). Here, we find that mouse somatosensory brain regions undergo stereotyped transformations in early postnatal development that expand the populations of neurons dedicated to glabrous skin. Our findings point to a model in which this remapping occurs at the first synapse between peripheral mechanoreceptors and their central targets in the brainstem.

### Determinants of body representation in the central nervous system

Multiple lines of evidence suggest that the central representation of touch is at least partly determined by skin innervation density (Lee and Woolsey, 1975; Corniani and Saal, 2020). Enrichment of sensory receptors in specific skin targets may reflect increased production of somatosensory neurons at the relevant axial levels, enhanced attraction of peripheral axonal projections, or decreased cell death of sensory neurons that innervate those skin regions, producing differences in innervation density. Synaptic expansion, the ratio of presynaptic neurons to postsynaptic neurons in a neural circuit, is determined in part by the degree of convergence in the system (Litwin-Kumar et al., 2017). As our fiber fraction measurements indicate systematic differences in convergence that depend on skin region, we propose that synaptic expansion at the LTMR-PN synapse in the brainstem acts multiplicatively with innervation density to emphasize skin regions with particular behavioral significance, such as the glabrous skin of the paws. In this view, different skin regions uniquely instruct expansion of mechanoreceptor synapses in the brainstem to shape a collective central representation. This model reconciles previously observed discrepancies between innervation density and central representation across the body in multiple species and quantitatively fits our data for mouse hindlimb and forelimb glabrous and hairy skin (Catania and Kaas, 1997; Corniani and Saal, 2020; Lee and Woolsey, 1975). We propose that skin region dependent patterns and strengths of LTMR synaptic connections within the brainstem reduce evolutionary constraints by allowing body maps to be flexibly tailored to disparate mammalian body forms.

Disproportionate representation of glabrous skin corresponds with greater tactile acuity on these skin surfaces and may underlie fine sensory-motor control needed for dexterous object manipulation. In the visual system, allometric scaling laws that relate the size of visual thalamus and cortex across primate species have led to the proposal that synaptic expansion enables higher order sensory areas to explicitly represent new features of a stimulus (i.e. direction of stimulus movement) while maintaining topography (Stevens, 2002). We predict that expanded glabrous skin representation in the brainstem may support the early, subcortical emergence of neural feature tuning. Parallel feature computations in early somatosensory circuits are proposed to increase the speed at which the nervous system can act on mechanical feedback (Tuthill and Wilson, 2016). Alternatively, the increased efficacy of synaptic transmission in glabrous skin tactile circuits of the brainstem might underlie reliable encoding of weak tactile stimuli, such as a subtle contour or edge of a held object. As the extent of disproportionate representation of glabrous skin in the brainstem is equivalent to that in somatosensory cortex, future investigations in brainstem might reveal the logic of disproportionate body representation more generally.

### The development of body representation

Functional measurements in pre-term and full-term human infants suggest that while somatotopic organization of cortical responses to touch is present at birth, these responses undergo rapid maturation in the first weeks of life (Allievi et al., 2016; Dall’Orso et al., 2018). Defining the developmental mechanisms that shape the neural representation of touch from the periphery to cortex will form a foundation for a neurophysiological understanding of the acquisition of the brain’s sense of the body and may inform our understanding of neurodevelopmental disorders in which somatosensation may be disrupted.

Peripherally-mediated activity plays a critical role in the development of the visual and auditory systems, and touch-evoked developmental activity has been proposed to be a determinant of central representation sizes in the somatosensory system (Catania, 2001). To our surprise, however, we found that animals lacking mechanically evoked activity in somatosensory neurons retained the relationship between skin target identity and the strength of LTMR synaptic outputs in the brainstem. While we cannot rule out that other aspects of LTMR development or connectivity are affected in *Piezo2* conditional mutants, our findings suggest that differential synaptic weights between glabrous and hairy skin mechanosensory neurons are established through mechanisms that are largely if not entirely independent of mechanically evoked activity.

Discoveries in the star-nosed mole have implicated the brainstem as a potential locus of disproportionate representation (Catania and Kaas, 1997; Catania et al., 2011). In this system, a pattern of cytochrome-oxidase densities in brainstem forms over development, with the behaviorally most-relied upon regions of the star appendage forming first and occupying a disproportionate area (Catania, 2001). It was proposed that disproportionate body representation might reflect a model in which the earliest axonal arbors to arrive to the brainstem have an advantage over later arrivals. This model is unlikely to account for the disproportionate representation of glabrous skin seen here, as innervation density is proportionate to central representation in juvenile mice, and differences in central representation emerge long after all projections from the periphery have reached the brainstem.

The mechanisms that determine the identity and function of the peripheral neurons whose endings are embedded in the skin are not fully understood. Our experiments in *Piezo2* mutants suggest that skin type-dependent synaptic expansion occurs in the absence of touch, and that tactile experience is not required to skew central body maps towards glabrous skin. Instead, we propose that the ventral paw/glabrous skin either i) provides an instructive signal to developing LTMRs such that those neurons innervating glabrous skin form more synaptic connections in the brainstem, or ii) generates cues that attract LTMRs that are pre-programmed to make more synaptic connections in the brainstem. Peripheral targets play key roles in controlling somatosensory neuron maturation, and skin type can instruct basic properties of peripheral neuron identity, such as the neurotransmitter they produce, suggesting a mechanism that could produce enlarged glabrous representations (Schotzinger and Landis, 1988; Luo et al., 2007). We extend these ideas by showing that skin terminal endings characteristic of both glabrous and hairy skin can be formed by the same neuron, and further propose that peripheral target identity also controls the strength of a neuron’s central synaptic connections, and ultimately, the proportion of neurons that represents the skin target. This mechanism may extend beyond the central representations of gentle touch to the skin, as a recent study reported that the central arbor morphologies of a subset of nociceptors also vary by body location (Olson et al., 2017).

The observation that neurons with peripheral arbors in glabrous-adjacent regions of hairy skin adopt glabrous skin central projection phenotypes suggests the existence of a diffusible factor or cue produced in glabrous skin that preferentially preserves synaptic connections in the brainstem during early postnatal periods of developmental refinement. A substantial body of literature demonstrates that somatotopy develops in an outward-to-inward fashion, and indicates that body maps established at lower levels of the somatosensory hierarchy drive the organization of higher regions (Iwasato and Erzurumlu, 2018). We propose that skin type influences the development of mechanoreceptor synaptic connectivity in the brainstem and is thus a powerful determinant of central touch representation.

## Acknowledgements

We thank Ofer Mazor and Pavel Gorelik from the HMS Research Instrumentation Core Facility for helping to design and fabricate the optical stimulator with support from NEI P30 Core Grant for Vision Research EY012196, Chinfei Chen for advice on fiber fraction electrophysiological measurements, Ardem Patopoutian for *Piezo2* mutant mice, Richard Born and Vladimir Berezovskii for assistance with attempts at viral labeling of LTMR brainstem synapses in nonhuman primate, Evan Feinberg and Daniel Dombeck for advice on the brainstem imaging preparation, David Paul for generating viruses used for anatomical experiments, Annie Chen, Sabrina Belozerova, Victoria Malarczyk, Alexandra Malarcyzk, and Connie Tsan for assistance with genotyping and mouse breeding, and Rachel Essner for assistance with multiphoton brainstem imaging. We thank Caiying Guo and the Gene Targeting and Transgenic Facility at the Janelia Research Campus of the Howard Hughes Medical Institute for generating mouse lines. We are grateful to Mark Andermann, Soha Ashrafi, Greg Bashaw, Chinfei Chen, Evan Feinberg, Annie Handler, Elizabeth Hong, Shan Meltzer, Aniqa Tasnim, John Tuthill, and Rachel Wilson for valuable comments on the manuscript. This work was supported by a William Randolph Hearst Fellowship (B.P.L), a Goldenson Fellowship (B.P.L), NIH grants F32 NS095631-01 (B.P.L.), F32-NS106807 (C.S.), DP1 MH125776 (C.D.H.), R01 NS089521 (C.D.H.), R01 NS97344 (D.D.G), an HMS Dean’s Innovation Grant in the Basic and Social Sciences (C.D.H. and D.D.G.), and the Edward R. and Anne G Lefler Center for Neurodegenerative Disorders (D.D.G.). D.D.G. is an Investigator of the Howard Hughes Medical Institute.

## Author Contributions

B. P.L., C.S., and D.D.G. conceived the study. B.P.L. performed multiphoton imaging and electrophysiological experiments with assistance from E.H., A.J.E, and S.R., and input from C. D.H.; C.S. performed anatomical, morphological, and synapse histology experiments and designed and characterized transgenic mice with the assistance of N.A., S.R., B.P.L., I.A., Y.Z., L. B., C.K., J.T.H. and A.R.M.; B.P.L., C.S., and D.D.G. wrote the paper with input from all authors.

## Lead Contact and Materials Availability

Further information and requests for resources should be directed to and will be fulfilled by David Ginty.

## Experimental Model and Subject Details

All mice used in the study are of mixed background. The dual recombinase-dependent *Tau^FSFiAP^* knock-in allele was generated at the Janelia Campus Research Gene Targeting and Transgenic Facility using standard ES cell targeting. Briefly, a FRT-2XSTOP-FRT-5’PLAP-loxP-3’PLAP-loxP-attB-Neo-attP cassette was introduced by homologous recombination into the *Tau (Mapt)* gene, replacing the first ATG of the Tau coding sequence. The PLAP coding sequence encodes human placental alkaline phosphatase and its 3’ half is inverted and flanked by head-to-head loxP sites, preventing expression in the absence of Cre recombination. *TauFSF^iAP^* heterozygous mice were generated by mating chimeric males to females constitutively expressing *PhiC31(R26-PhiC3lo*, JAX #007670) to remove the neomycin selection cassette. Successful excision of the Neo cassette was confirmed in the next generation by PCR genotyping. The following primers were used to genotype *TauFSF^iAP^* animals for the presence of the reporter allele:

forward 5’-GGATGGGAAACTGAGGCTCT-3’; reverse 5’ ATGGTGGCGAATTCCAAATCA 3’ *R26_FSF-ReaChR_* mice were generated from *R26_FSF-LSL-ReaChR_* (JAX # 024846)(Hooks et al., 2015) mice through germline excision of the loxP-flanked STOP cassette. Previously published mice used in this study include: *Scnn1a-Tg3-Cre* (JAX #009613), *Ai94(TITL-GCaMP6s)-D;CaMK2a-tTA* (JAX # 024115)(Madisen et al., 2015), *Brn3a_CKOAP_* (Badea et al., 2009a), *R26_IAP_* (Badea et al., 2009b), *Gad2_T2A-NLS-mCherry_* (JAX#023140)(Peron et al., 2015), *Cdx2-Cre* (Coutaud and Pilon, 2013), *Advillin_FlpO_* (Choi et al., 2020), *Rosa26_FSF-LSL-tdTomato_* (Ai65) (JAX#021875)(Madisen et al., 2015), *Rosa26_LSL-synaptophysin-tdTomato_* (Ai34) (JAX#012570), _Ret_CreER__ (Luo et al., 2009), _TrkB_CreER__ (Rutlin et al., 2014), _TrkC_CreER__ (Bai et al., 2015), _Ptgfr_CreER__ (this line will be characterized elsewhere), _Pvalb_T2A-FlpO__ (JAX # 022730)(Madisen et al., 2015), *Piezo2_-_* (Nonomura et al., 2017), and *Piezo2_fl_* (JAX #027720)(Woo et al., 2014).

Mice were handled and housed in standard cages accordance with the Harvard Medical School and IACUC guidelines. Both male and female mice were used in all experiments.

## Method Details

### Acute Brainstem Slice Recordings

Animals were transcardially perfused with ice-cold choline solution comprising (in mM) 92 choline chloride, 2.5 KCl, 1.2 NaH2PO4, 30 NaHCO3, 20 HEPES, 25 Glucose, 5 sodium ascorbate, 2 thiourea, 3 sodium pyruvate, 10 MgSO_4_, and 0.5 CaCl_2_ adjusted to pH 7.3 with Tris Base. The brainstem and rostral spinal cord were dissected away, immersed in ice-cold choline solution and trimmed at the pons rostrally and the lower cervical spinal cord caudally. Tissue was then mounted dorsal side up and adjusted such that the surface of the dorsal column nuclei was aligned with the direction of the vibratome blade (VT1200S, Leica). An initial 25 micron slice through the remaining dura was made to permit electrode access. Subsequently, a single horizontal slice (280 – 300 μm thick) was cut such that it included the dorsal column nuclei and rostral spinal cord. Slices were transferred to an incubation chamber and bathed in ACSF comprising (in mM) 125 NaCl, 2.5 KCl, 2 CaCl2, 1 MgCl2, 26 NaHCO3, 1.25 NaH2PO4, and 25 glucose, bubbled with 95% O2/5% CO2 for 40-60 minutes prior to recording. Immediately prior to placing them in the recording chamber, slices were allowed to adhere to poly-L-lysine coated coverslips (15mm, Warner Instruments) and placed with their dorsal surface exposed for whole-cell recording.

Whole-cell recordings were obtained with an internal solution containing (in mM): 135 Cs-methanesulfonate, 10 Hepes, 1 EGTA, 3.3 QX314 (Cl-salt), 4 Mg-ATP, 0.3 Na-GTP, 8 Na-phosphocreatine, 5 BAPTA, and 10 neurobiotin citrate, adjusted to pH 7.3 with CsOH. Wholecell current-clamp recordings used an internal solution contained the following (in mM): 100 K-methanesulfonate,11.8 NaCl, 1.8 MgCl2, 3.6 Mg-ATP, 0.45 Na-GTP, 12.7 phosphocreatine, 9.1 EGTA, 0.9 CaCl2, 9 HEPES, and 14 KOH. Reported voltages are corrected for a −8 mV liquid junction potential between the K-methanesulfonate solution and the ACSF.

### Fiber fraction measurements

Mice were anesthetized with isoflurane and placed in a small animal stereotaxic frame (David Kopf Instruments). Under aseptic conditions, the skull was exposed and two craniotomies centered at 1.5 mm posterior and 1.9 mm lateral (bilateral) from bregma were created. To retrogradely label thalamic projection neurons residing in the DCN for whole cell recordings, a borosilicate glass pipette was lowered into the thalamus to a depth of 3.1 mm, allowed to rest for 1 minute, then retracted to 3.0mm. After 5 minutes, 100-200 nL of fluorescent retrobeads (Lumafluor) were injected at a rate of 100 nl·min-1 using a Microinject system (World Precision Instruments). During surgery, mice received buprenorphine and carprofen before being returned to their home cage for at least 2 days prior to slice recording.

Whole-cell patch clamp recordings were obtained from fluorescently labeled thalamic projection neurons in the gracile nucleus under visual guidance. Voltage-clamped currents were recorded with an Axopatch 700A amplifier, low-pass filtered at 10 kHz, and sampled at 40kHz by a 16 bit A/D converter (USB-6343, National Instruments) and acquired in Wavesurfer (Janelia). Neurons were clamped at −70 mV and estimates of access resistance based on the height of the fast current transient during test voltage steps were 5-20 MOhm. The dorsal column was stimulated electrically with a steel bipolar electrode (30204, FHC) or optically in mice expressing ReaChR in glabrous or hairy skin innervated Aβ-LTMRs. To optogenetically stimulate axons, the objective was positioned 1-4 millimeters caudal to the gracile nucleus and the field stop adjusted to produce a ~100 micron spot of light on the ipsilateral dorsal column. Brief pulses (100-700 microsecond) of light from an LED light source (M470F3, Thor Labs) was delivered and the intensity adjusted to attain successes and failures in roughly equal proportion, thus stimulating a single fiber. Heterotypic inverse fiber fraction measurements are expressed as the ratio of the amplitude of the current evoked by saturating electrical stimulation to the optogenetic single fiber amplitude. Homotypic inverse fiber fraction measurements are expressed as the ratio of the amplitude of the current evoked by saturating optical stimulation to the optogenetic single fiber amplitude (Litvina and Chen, 2017).

### Stereotactic injections for brainstem calcium imaging

Brainstem projection neurons were retrogradely transduced through injection of AAV2-retro viruses to their brain targets. Three to four weeks prior to imaging of the adult brainstem (or one week prior for P10-P14 imaging), mice were injected with sustained release buprenorphine (0.1 mg/kg), anesthetized with isoflurane and placed in a small animal stereotaxic frame (David Kopf Instruments). To target thalamic projection neurons from brainstem, the skull was exposed under aseptic conditions and two craniotomies centered at 1.5 mm posterior and 1.9 mm lateral (bilateral) from bregma were created. A borosilicate glass pipette was lowered to a depth of 3.2 mm, allowed to rest for 1 minute, then retracted to 3.1mm. After 5 minutes, 100-200 nL of rAAV2-retro jGCaMP7s (~1e13 gc/mL, Addgene) was injected at a rate of 70 nL·min-1 using a Microinject system (World Precision Instruments). This procedure was repeated a second time to target a different location along the AP axis with a second injection in the coordinate range 1.3-1.7mm posterior to bregma. To target neurons that project to the inferior colliculus, the skin was retracted under aseptic conditions and craniotomies placed bilaterally at .5mm posterior to lambda and 1.5 mm lateral to the midline. A glass pipette was lowered to a depth of 1mm from the surface of the brain, then retracted to 0.7mm. After 2 minutes, 200 nL of rAAV2-retro jGCaMP7s (~1e13 gc/mL, Addgene) was injected at a rate of 70 nL·min-1. This procedure was repeated at a second location along the A-P axis.

### *In vivo* brainstem imaging surgical preparation

Mice were anesthetized with urethane (0.5-1.5 mg/g) and chlorprothixene (5 mg/kg) and placed on a bite bar (Kopf Instruments) on a custom heated recording platform. The head was flexed downward at a 45 degree angle and an incision made in the skin and overlying muscle to expose the cervical paraspinal muscles. Next, the paraspinal muscles were retracted laterally, exposing the skull and first cervical vertebra. A hemicircular cut in the skull was made with a dental drill and a posterior section of the skull overlying the cerebellum removed. The posterior atlanto-occipital membrane was carefully removed through incision directly rostral to the atlas and laterally to the occipital condyles. The dural membrane was then retracted caudally through insertion of a Bonn microprobe (Fine Science Tools) between the cerebellum and dorsal surface of the brainstem. An imaging chamber that stabilizes the brainstem and displaces the cerebellum rostrally was fabricated from a circular 3mm glass coverslip (CS-3R, Warner Instruments) and flanged stainless tube (Microgroup) attached to a titanium plate and mounted to a 3d micromanipulator (MMN-1, Narishige). The assembly was lowered onto the caudal brainstem just rostral to the atlas and downward pressure applied to the brainstem surface until there was no apparent motion. The assembly was then translated dorsally to displace the cerebellum, centered on the obex, and the interface between the imaging chamber and skin sealed with adhesive. Healthy long-term preparations were associated with pressures that stabilized the brainstem cardiac and respiratory motions while avoiding full exsanguination of the spinal vertebral and penetrating arteries. In a small fraction of animals, arterial elaboration over the gracile nucleus made imaging infeasible.

### Multiphoton calcium imaging, mechanical stimulus, and analysis

Two-photon calcium imaging was performed using a resonant multiphoton microscope (Scientifica) equipped with a piezoelectric objective Z-stepper (P-726, Physik Instrumente). All images were acquired using a 16X, 0.80 NA, 3 mm WD objective (Nikon) with scan mirror excursions that produced a 510 x 510 μm^2^ field of view. Volumetric data was captured by acquiring 4-6 planes with a sawtooth scan configuration, yielding a volumetric imaging rate of 7.5 – 5 Hz. Each FOV was at least 20 μm below or above an adjacent FOV for cell body imaging. Excitation light was provided by a Ti:Sapphire laser (80 MHz, Vision S, Coherent) tuned to 910-925nm with pre-chirp dispersion compensation. Laser powers measured at the front aperture of the objective were between 15 – 60 mW depending on imaging depth. Complete coverage of the gracile nucleus typically required eight tiled multiphoton imaging volumes.

Calcium imaging time series (512 x 512 pixels) were initially registered to a reference image to correct for motion in the rostal-caudal and medial-lateral directions. ROIs containing putative cells were automatically detected based on their morphology and time-varying indicator fluorescence from the aligned image series and manually curated to remove extraneous cell profiles. Signals were extracted as the average value in each ROI sampled at the volumetric frame rate using Suite2P and corrected for signal contamination from surrounding neuropil as F_neuropil_corrected_(t) Froi(t) – .7 x F_neuropil_(t). Touch stimuli were ~10mm long skin strokes delivered once per second from the proximal to distal direction with a soft, 1.5 mm wide brush (Blick Art Materials, Boston MA). Stroke was delivered for 8 seconds to a given body region with 16 second inter-trial intervals. Cells were considered responsive if their average fluorescence exceeded the baseline plus 2.5 times the standard deviation of the baseline for 30% of the stimulus period.

### Dorsal Column Injections

For dorsal column injections, mice were placed on the stereotactic frame with their neck bent at a 45 degree angle. The dorsal column was exposed between the C1-C2 levels through retraction of the paraspinal muscles and the overlying laminar and dural membranes removed prior to injection. For neuronal innervation density estimates, 100-200 nL of rAAV2/1-Cre virus (Penn Viral Core, diluted to 2E12 vg/mL) was injected into the dorsal column 200 microns below the spinal cord surface. For optogenetic measures of convergence, 300 nL of undiluted virus (1e13 gc/mL) was injected bilaterally. Incisions were sutured and the animal was allowed to recover before being returned to its home cage for postoperative monitoring.

### Skin Injections

Mice P6 and older were anesthetized with continuous inhalation of 2% isoflurane from a precision vaporizer for the duration of the procedure (5-10 min). The animal’s breathing rate was monitored throughout the procedure and the anesthetic dose adjusted as needed. The skin region of interest was swabbed with ethanol. Injections were done with a beveled borosilicate glass pipette. Forceps were used to stabilize the skin while the needle was inserted into the dermis, injecting 0.5-2 μL of virus. Mice were injected in 1-6 locations within the skin region of interest, depending on skin identity and the amount of labeling desired. Mice harboring CreER alleles along with a dual recombinase fluorescent or alkaline phosphatase reporter were injected with AAV2/1-hSyn-FlpO (1.37e13, Boston Children’s Hospital Viral Core) to target select LTMR subtypes. Mice harboring a Cre dependent reporter allele were injected with AAV2-retro-hSyn-Cre (1.2e13, Addgene 105553), while mice harboring a Flp-dependent reporter allele were injected with AAV2/1-hSyn-FlpO (1.37e13, Boston Children’s Hospital Viral Core). Wild type mice were injected with AAV2/1-hSyn-synaptophysin-TdTomato (1e14, UPenn Viral Core) or AAV2/1-synaptophysin-mScarlet (1.4e13, Boston Children’s Hospital Viral Core). For all injections, a small amount of fast green (Sigma F7252-5G) in 0.9% saline was added to the virus. After injection, mice recovered from anesthesia on a warm pad while being monitored. Mice were returned to the litter once normal activity had resumed. Five weeks post-injection (1 week post-injection for p14 animals) animals were sacrificed by transcardial perfusion under isoflurane anesthesia.

### Perfusions

Mice were anesthetized under isoflurane and transcardially perfused with 5-10mL of Ames Media (Sigma) in 1xPBS with heparin (10 U/mL), followed by 5-10 mL of 4% paraformaldehyde (PFA) in 1xPBS. After perfusion, the skull, vertebral column, and central nervous system were removed and post-fixed in 4% PFA in 1xPBS at 4°C overnight. Skin samples were removed and treated with commercial depilatory cream (NAIR, Church and Dwight Co.; Princeton, NJ) for 3-5 minutes, then washed with soap and water. Skin samples were fixed overnight in Zamboni’s fixation buffer with gentle agitation at 4°C. Samples were washed 3×10min in 1xPBS at room temp after overnight fixation before fine dissection.

### Immunohistochemistry of free-floating sections

The brain, brainstem, and spinal cord were removed from the skull and vertebral column. The skull was cut down the midline and pulled laterally to remove. The dorsal and ventral sides of the vertebral column were removed. Then, forceps were used to remove the dorsal root ganglia (DRG) bilaterally from the vertebral column and surrounding tissue, retaining the connection to the spinal cord. This was done for all DRGs working from lumbar to cervical levels. Once all DRGs had been removed, the brain, brainstem, and spinal cord could be extracted from the vertebral column and stored in 1x PBS.

Sectioning of tissue samples was done using a vibrating blade microtome (Leica VT100S). Samples were embedded in 3% agarose in 1x PBS. Once solidified, samples were trimmed and attached to the sectioning plate with superglue. Transverse brainstem and spinal cord sections were taken at 100 μm while horizontal brain sections were taken at 200 μm. Tissue samples were then rinsed in 50% ethanol/water solution for 30 minutes, followed by 3x 10 min in 2xPBS. Tissue samples were incubated in a mixture of primary antibodies in high salt Phosphate Buffer Saline (2x PBS) containing 0.1% Triton X-1000 (2xPBSt) for 72 hours at 4°C. Primary antibodies used in this study: Guinea pig anti-Vglut1 1:1000 (Sigma AB5905), Rabbit anti-Homer1 1:500 (CedarLane 160003SY), Goat anti-mCherry 1:500 (CedarLane AB0040-200), Chicken anti-GFP 1:500 (Aves GFP-1020), Rabbit anti-tRFP 1:500 (Evrogen AB233), Rabbit anti-S100 1:300 (VWR/ProteinTech 15146-1-AP), Chick anti-Neurofilament 200 kDa 1:500 (Aves NFH). The tissue was rinsed 3×10 min with 2xPBS before incubation in the secondary antibody solution for 24 hours at 4°C. The secondary antibody solution was made up in 2xPBSt and contained species-specific Alexa Fluor 488, 546, and 647 conjugated IgGs. Secondary antibodies used in this study: Donkey anti-Guinea pig Alexa Fluor 647 1:500 (Jackson Immunoresearch 706-605-148), Donkey anti-Rabbit Alexa Fluor 488 1:1000 (ThermoFisher A-21206), Donkey anti-Goat IgG Alexa Fluor 546 1:1000 (LifeTech A11056), Donkey anti-Chicken IgG Alexa Fluor 488 1:1000 (Jackson Immunoresearch 703-545-155), Donkey anti-Rabbit IgG DyLight 405 1:1000 (Jackson Immunoresearch 711-475-152). The tissue was then mounted on glass slides, coverslipped in VectaMount mounting medium (Vector labs H-5000) and stored at 4°C until imaging.

### Immunohistochemistry of skin cryosections

Paws were roughly dissected from perfused mice and hair was removed with a commercial depilatory cream (Nair, Church & Dwight). Glabrous and hairy tissue was finely dissected from the paws, cryoprotected in 30% sucrose in 1x PBS at 4**°**C for 2 days, embedded in OCT (1437365, Fisher), frozen using dry ice, and kept at −80**°**C. The tissues were cyrosectioned (25 μm) using a cryostat (Leica) and collected on glass slides (12-550-15, Fisher). Sections were washed 3 times for 5 minutes each with 1x PBS containing 0.1% Triton X-100 (0.1% PBST), incubated with blocking solutions (0.1% PBST containing 5% normal goat serum (S-1000, Vector Labs) or normal donkey serum (005-000-121, Jackson Immuno)) for 1 hour at RT, incubated with primary antibodies diluted in blocking solutions at 4**°**C overnight, washed 3 times for 10 minutes each with 0.1% PBST, incubated with secondary antibodies diluted in blocking solutions at 4**°**C overnight, washed again 4 times for 10 minutes each with 0.1% PBST (DAPI solution was included in the second wash at 1:5000 dilution), and mounted in Fluoromount-G mounting medium (Fisher 0100-01).

### Multielectrode Array (MEA) Recordings

MEAs were inserted into the L4 dorsal root ganglia (H6b-32ch, Cambridge Neurotech), gracile nucleus of the brainstem (E1-16ch, Cambridge Neurotech), primary somatosensory cortex (F3-32ch) and inferior colliculus (H6b-32ch, Cambridge Neurotech) with a 3-axes micromanipulator (IVM mini, Scientifica). MEA signals were high-pass filtered at 200 Hz, amplified (RHD2132) and acquired at 20kHz (RHD2000, Intantech) for offline processing.

### Spike detection and cluster assignment

Data was converted into contiguous binary files (https://github.com/peltonen/kwik-tools) and processed using JRClust (Janelia). After pre-processing and filtering, spikes that exceeded detection thresholds defined as qq factors (RQ Quiroga, 2004) that were adjusted (4.5-6.0) based on recording site and probe. Spikes were detected and assigned to clusters corresponding to individual or multiples units with JRClust (https://github.com/JaneliaSciComp/JRCLUST). These initial clusters were manually curated to remove artifacts that did not correspond to spike waveforms, and clusters were examined and split if their PCA decompositions were not monodisperse. This collection of clusters was then examined and merge operations performed if consideration of the average spike waveform and spiking cross-correlation indicated that two clusters corresponed to the same neuron. Manual curatation resulted in putative isolated single units with intra-cluster waveform correlations greater than 0.9, inter-cluster waveform correlations less than 0.1, and a 20-fold depletion of spiking events at t=0 msec in the cross-correlogram, and clusters that did not meet this criteria were not considered in the data set. Clusters computed in this manner contained units with spatially distinct receptive fields, suggesting that units were drawn from non-overlapping sources. Spikes event times for each cluster were exported and processed in Python.

### Optical Skin Stimulation

Hair was removed from the hindlimb ipsilateral to site of optical stimulation with a commercial depilatory cream (Nair, Church & Dwight). The leg and foot were immobilized with adhesive tape and positioned underneath a custom-built optical skin stimulator. Next, an image of the animal was taken along the axis of optical stimulation through a high-pass dichroic with a monochrome camera (FLIR Systems). The stimulator mirrors were parked at their origin and the skin illuminated with low power laser light. A subsequent image of the paw and laser illuminated spot was captured. This procedure allowed for the centering and orienting of maps derived from optical stimuli onto the skin.

The optical stimulator was constructed from a laser light source (445 nm, 100 mW), a beam expander (GBE03-A, Thor Labs), galvometer scan mirrors (6210H, Cambridge Technologies), an f-theta lens (FTH-160-1064-M39, Thor Labs), and a front end dichroic (FF660-Di02-25×36 Semrock) for dividing the optical path between the stimulator and camera.

### Optical Receptive Field Determination

A spike-triggered average based analysis was used to describe the receptive fields of mechanically sensitive neurons rendered optically responsive through expression of the channelrhodopsin variant ReaChR (Hooks et al., 2015). Optical receptive fields were estimated for each recorded unit that responded to sparse white noise optical stimulation of the skin according to the equation (cite Pillow):

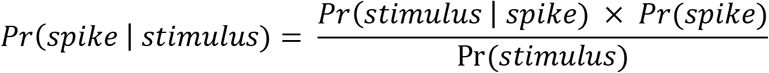

To compute Pr(spike | stimulus), we first binned spikes from each unit at 1 msec to produce a [N x 1] binary vector of spike occurrences, where N is the number of discrete time bins encompassing the recording. Next, the stimulus was binned at each time step into a 2d spatial histogram with 100 micron bin edges to produce a matrix indicating stimulation of a given location. This was subsequently flattened in column-major order to produce a [M x N] stimulus matrix, where M is the number of (binned) spatial locations and N is the number of discrete time bins. Next, a design matrix [(M x mkt) x N] containing the relevant regressors for each time bin was constructed from the stimulus matrix by collecting all space-time stimulus elements within a time window of nkt samples (Pillow). The spike triggered stimulus average with dimensions [M x nkt] is the product of the design matrix and the spike vector. This was used to compute the quantity Pr(spike | stimulus) according to the equation above by dividing by the raw stimulus ensemble.

### Cortical windows and functional imaging of somatosensory cortex

A circular craniotomy with a diameter of 3.1 mm was made over right primary somatosensory cortex (stereotaxic coordinates: 0 mm anterior-posterior, 1.8 mm lateral of bregma). A glass plug consisting of a single 5 mm diameter coverslip on top of two 3 mm diameter coverslips (#1 thickness; CS-5R and CS-3R, Warner Instruments) were assembled using UV-curable optically transparent adhesive (Norland Optics) and cemented to the skull using Metabond applied to the perimeter of the 5 mm coverslip. A titanium ring was mounted on top of the headplate and connected to the microscope’s objective lens through a cylinder of black rubber to prevent light contamination. The mouse was placed on a heated imaging and immobilized such that the paws on the side contralateral to the imaging window were extended for mechanical stimulation.

### Whole-mount alkaline phosphatase staining of the skin, spinal cord, and brainstem for sparse neuronal labeling

To estimate relative peripheral innervation densities across different body regions, *Brn3a^cKOAP^/+* mice aged 4-8 weeks (for adult time points) or 7 days (for P14 timepoint) were stereotactically injected with AAV2/retro-Cre virus into the dorsal column at cervical levels C1-C2 as described above. Four weeks post virus injection (1 week for p14 animals), animals were transcardially perfused using 4% paraformaldehyde. Skin and paws were cut off, hair was removed using NAIR, and skin was post-fixed in Zamboni solution at 4°C overnight. Spinal cords and brainstems were post-fixed in 4% PFA, 4°C overnight. The next day, tissue was washed three times in 1xPBS and stored in PBS at 4°C until further processing.

To visualize brainstem arbors by sparse labeling, three different strategies were used. To sparsely label all cutaneous neurons that send ascending projections to the dorsal column nuclei, *Brn3a^cKOAP^* animals aged 8-15 days were injected with AAV2/retro-hSyn-Cre virus into hairy or glabrous paw skin. 4-5 weeks post virus injection, animals were transcardially perfused with 4% paraformaldehyde, and skin and CNS were collected as described above. After fine dissection, brainstems were processed for whole-mount alkaline phosphatase staining (see below). For all animals with sparse labeling of DCN arbors in the gracile or cuneate nucleus (1-2 arbors per animal), the corresponding paws were dissected and stained to identify the position and morphology of the labeled cutaneous arbor. Only neurons with somatotopically unambiguously identified peripheral arbors were used for further analysis. To label specific genetic classes of LTMRs, *Ret^CreER^* or *Ptgfr^CreER^* mice were crossed to *Brn3a^cKOAP^* mice to generate *CreER/+; AP/+* animals. Low dosage tamoxifen (~0.01 mg) was delivered by oral gavage of pregnant females at E11.5 (*Ret^CreER^*) or intraperitoneal injection of pups at P21 (*Ptgfr^CreER^*). This method yielded a low success rate of animals with unambiguous, sparse labeling in paws. Therefore, to allow for higher throughput sparse labeling of genetically identified arbors innervating specific skin regions, we used *TauFSFiAP*, a newly generated dual Cre and FlpO-dependent AP allele. Animals containing CreER alleles (*Ret^CreER^, TrkC^CreER^*, or *Ptgfr^CreER^*) and the *Tau^FSFiAP^* allele were administered high doses of tamoxifen to maximize Cre-dependent recombination. Then, AAV-FlpO virus was injected to glabrous or hairy paw skin at P8-P10. 5-6 weeks post virus injection, animals were transcardially perfused with 4% paraformaldehyde and their brainstems and paws were dissected and processed for whole-mount alkaline phosphatase staining. The *TauFSFiAP* allele was also used to sparsely label neurons innervating Meissner corpuscles by generating *Ret^CreER^/+; Pvalb^FlpO^/+; TauFSF^iAP^/+* mice and delivering 0.025-0.05 mg tamoxifen intraperitoneally at P5-P7.

Whole-mount alkaline phosphatase staining was performed as previously described (Liu et al., 2007). Briefly, after post-fixation, tissue was washed 3x in 1XPBS, incubated at 65°C for 2-2.5 hours, and washed with B3 buffer (0.1 M Tris pH 9.5, 0.1 M NaCl, 50 mM MgCl2, 0.1% Tween-20) 3×5 min, at room temperature. AP signal was detected by incubating tissue in NBT/BCIP (Roche) diluted at 3.4 μg/mL in B3 buffer at room temperature, for 22-30 hrs with gentle rocking. To stop the AP reaction, tissue was incubated in 4% PFA at room temperature for 1 hour, washed 3x 5 min in 1XPBS, and then dehydrated (1 hour 50% ethanol, 1 hour 70% ethanol, 1 hour 100% ethanol followed by overnight incubation in 100% ethanol) before clearing with BABB (1 part benzyl benzoate:2 parts benzyl alcohol) for 4 hours at room temperature. Paws and brainstems were imaged on a Zeiss AxioZoom microscope.

### Quantification of pre-synaptic boutons in the GN/CN

To label pre-synaptic boutons of cutaneous sensory neurons, *R26^syt-tdTomato^* (Ai34) were injected with AAV2/retro-Cre virus, or wild type animals (CD1) were injected with AAV2/retro-synaptophysin-tdTomato or AAV2/retro-synaptophysin-mScarlet virus into the hairy or glabrous paw dermis at P5-P15. Animals were transcardially perfused with 4% paraformaldehyde 5-6 weeks post injection for adult time points (P50-P70), or 1 week post injection for developmental time points (P14). After overnight post-fixation in 4% PFA at 4°C and three washes in 1xPBS, dorsal column nuclei were finely dissected and sectioned transversely on the vibratome at 90-100 μm/section. Antibody staining was performed as described above. Sections were mounted in VectaMount and imaged on a Zeiss LSM 700 confocal microscope using a 40x oil-immersion lens. To quantify boutons, syn-FP+ puncta were identified semi-automatically using Imaris. Automatic thresholding and background subtraction were applied to all images. Total syn-FP+ puncta were identified using the Spot tool with manual curation. Co-localization with vGlut1 or Homer1 puncta was assessed manually after bouton identification for a subset of images.

### Dual Retrograde Tracing

AAVs encoding nuclear restricted fluorescent proteins, rAAV2/retro-hSyn-H2B-GFP-bGH (5.3E12 gc/mL) and AAV2/retro-hSyn-H2B-mTagBFP2-bGH (3.17E13 gc/mL), were stereotactically injected into the thalamus and the inferior colliculus (approximately 300nl per location) of *Gad2^T2A-NLS-mCherry^* ‘Sections were imaged on a Zeiss LSM 700 confocal microscope. To quantify overlap of labeled nuclei, fluorescent nuclei were identified semi-automatically using Imaris. Quantification of labeled nuclei and colocalization was obtained using the Spot tool with manual curation.

### Tamoxifen treatment

Tamoxifen (Sigma T5648) was dissolved in 100% ethanol to 20 mg/mL, (30 minutes vortexing at room temperature) and then mixed in 2X volume sunflower seed oil (Sigma S5007) and vacuum centrifuged for 30 minutes to evaporate the ethanol. 20 mg/mL tamoxifen aliquots in sunflower seed oil were stored at −80°C until the day of use, when they were thawed 30 minutes at room temperature, while protected from light, and if necessary further dissolved in sunflower seed oil before administering to mice. Pregnant mice received 0.02 mg – 3 mg tamoxifen by oral gavage at E11.5 (to label Aβ RA-LTMRs using *Ret^CreER^* mice), E12.5 (to label Aβ SA1-LTMRs using *TrkC^CreER^* mice), or E13.5-E16.5 (to label glabrous skin innervating Aβ RA-LTMRs using *TrkB^CreER^* mice). To label Aβ SA1-LTMRs and Aβ field/free-LTMRs, *Ptgfr^CreER^* animals were administered tamoxifen postnatally by intraperitoneal injection at P21. The amount of tamoxifen delivered was titrated according to experimental needs.

### *In situ* hybridization

Detection of Piezo2 and Nefh transcripts was performed by fluorescent *in situ* hybridization, as described in (Sharma et al., 2020). Briefly, individual DRG ganglia were rapidly dissected from euthanized mice and frozen in dry-ice cooled 2 methylbutane and stored at −80°C until further processing. DRGs were cryosectioned at a thickness of 20 μm and RNA was detected using RNAscope (Advanced Cell Diagnostics) according to the manufacturer’s protocol. The following probes were used: Mm Piezo2 exons 43-45 (Cat# 439971-C3), tdTomato (Cat# 317041), Mm-Nefh (Cat# 443671). Sections were mounted in FluoroMount-G (Fisher 0100-01) and imaged on a Zeiss LSM 700 confocal microscope using a 40x oil-immersion lens.

### Quantification and Statistical Analysis

Statistical tests were conducted in Python using the SciPy stats module or in GraphPad Prism. Non-parametric tests and parametric tests were used for comparing two independent groups (Mann-Whitney-Wilcoxon test or Welch’s t-test), two related groups (Wilcoxon signed-rank test), and multiple groups (Kruskal-Wallis test with Bonferroni correction). p < 0.05 was considered significant. Additional details on sample sizes and statistical tests for each experiment can be found in figure legends and the main text.

