## Supplemental figures and legends for "Mechanoreceptor synapses in the brainstem shape the central representation of touch"

**Figure S1. Morphological and electrophysiological characterization of LTMRs on glabrous and hairy skin**

**A**. Characterization of Aβ LTMR responses to stroke on the glabrous or hairy skin. (top) Example cell-attached recordings of touch sensitive neurons with large cell bodies in DRG retrogradely labeled by CTB-647. (bottom) Number of spikes evoked by stroking glabrous or hairy skin across a population of Aβ LTMRs.

**B**. Innervation density at different proximal-distal positions of the body in $Brn3a^{cKOAP}$ mice injected with AAV2-retro-Cre in the dorsal column or dorsal column nuclei, normalized in each mouse to endings/square mm in trunk hairy skin.

**C**. The total surface areas of the peripheral arbors in the skin of sparsely labeled Aβ-LTMRs do not significantly change between P14 and adulthood (Welch's t-test). Areas were measured by tracing the convex hull of each arbor using Fiji.

**Figure S2. Principal neurons of the gracile and cuneate nuclei**

**A**. Strategy for visualizing cell bodies of neurons that project to the ventroposterolateral thalamus and shell of the inferior colliculus. An AAV2-retro-hSyn-H2B-GFP virus was injected into the thalamus and an AAV2-retro-hSyn-H2B-mTagBFP2 virus was injected into the inferior colliculus in Gad2-T2A-nls-mCherry mice.

**B**. Horizontal section of the brainstem at the level of the dorsal column nuclei, which comprises the external cuneate nucleus (ECN), the cuneate nucleus (CN), and the gracile nucleus (GN). Thalamic projection neuron, inferior collicular projection neuron, and Gad2+ cell bodies are found throughout the CN and GN but excluded from the ECN.

**C**. Higher magnification view of the GN of (A) and (B), showing that these three populations of neurons are largely non-overlapping.

**D**. Quantification of the degree of overlap compiled from two mice.

**E**. Circuit diagram showing the connections between the dorsal root ganglia (DRG), gracile/cuneate nuclei, ventroposterolateral (VPL) thalamus, inferior colliculus, and primary somatosensory cortex (S1) in the somatosensory hierarchy.

**Figure S3. Morphological characterization of the central arbors formed by Aβ-LTMRs that innervate glabrous or hairy skin**

**A**. The total length of axon collaterals of sparsely labeled Aβ-LTMRs in the dorsal column nuclei was quantified using the Simple Neurite Tracer in Fiji. Neurons innervating glabrous skin or the edge of paw adjacent to hairy and glabrous skin form brainstem axon collaterals of a greater total length than neurons innervating trunk hairy skin or back of paw hairy skin (**$p<0.001$ or *$p<0.01$ by Welch's t-test).

**B**. Length of axon collaterals in Aβ-LTMR arbors in the GN or CN plotted versus distance of each neuron's peripheral arbor to the edge of paw.

**C**. (left) Number of end points in the dorsal column nuclei formed by single Aβ-LTMRs that innervate different regions of paw, color-coded by LTMR subtype (same data as in Figure 3C, but color-coded by neuronal identity) **$p<0.001$ or *$p<0.01$ by Welch's t-test.

(right) Total length of axon collaterals formed by single Aβ-LTMRs that innervate different regions of paw, color-coded by LTMR subtype (same data as in panel A, but color-coded by neuronal identity). **$p<0.001$ or *$p<0.01$ by Welch's t-test.

**D**. (left) Number of end points in the dorsal column nuclei formed by single Aβ-LTMRs that innervate paw hairy skin, plotted against the size of the peripheral arbor formed by each neuron.

(right) Total length of axon collaterals in the dorsal column nuclei formed by single Aβ-LTMRs that innervate paw hairy skin, plotted against the size of the peripheral arbor formed by each neuron. No relationship was detected between the size of peripheral arbors and the number of end points or length of axon collaterals formed by their central arbors ($p>0.05$, simple linear regression).

**E**. Same as D, but for Aβ-LTMRs that innervate glabrous paw skin. No relationship was detected between the size of peripheral arbors and the number of end points or length of axon collaterals formed by their central arbors ($p>0.05$, simple linear regression).

**F**. Representative examples of the reconstructed central projections of Aβ-LTMRs in the gracile nucleus at different developmental stages.

**G**. Quantification of the number of end points in the dorsal column nuclei formed by Aβ-LTMRs that innervate hairy skin or trunk, at P5-P10 or in adult animals. At early postnatal stages, Aβ-

LTMRs that innervate hairy paw or trunk-level dermatomes form more complex axonal arbors in the gracile or cuneate nuclei than Aβ LTMRs of adult animals innervating similar skin targets ($p=0.001$, Welch's t-test).

**Figure S4. Additional bouton immunohistochemistry and functional imaging of thalamic projection neurons in the gracile nucleus over development**

**A**. Percentage of Aβ-LTMR syn-FP+ boutons that are apposed to Homer1+ puncta in the dorsal column nuclei. Each datapoint indicates boutons quantified in one animal. No difference was detected between Aβ-LTMRs innervating hairy and glabrous skin ($p>0.1$, Welch's t-test).

**B**. Percentage of Aβ-LTMR syn-FP+ boutons that co-localize with vGlut1+ puncta in the dorsal column nuclei. Each datapoint indicates boutons quantified in one animal. No difference was detected between Aβ-LTMRs innervating hairy and glabrous skin ($p>0.1$, Welch's t-test).

**C**. Percentage of Aβ-LTMR syn-FP+ boutons found at the end points of branches in the dorsal column nuclei. Each datapoint indicates boutons quantified in one animal. No difference was detected between Aβ-LTMRs innervating hairy and glabrous skin ($p>0.1$, Welch's t-test).

**D**. Profiles of P14 (left) and adult (right) thalamic projection neurons in the gracile nucleus that show significant responses to touch on hindpaw glabrous (red) or hindpaw hairy (blue) skin. Responses indicated exhibited calcium indicator fluorescence that exceeded three times the standard deviation of the baseline for at least a third of the stimulation window.

**E**. The percentage of thalamic projection neurons with multiregional receptive fields in the gracile nucleus, defined as having a significant response to touch at two or more regions on glabrous hindpaw skin, hairy hindpaw skin, thigh skin, or back skin declines over development (Kruskal-Wallis, Mann Whitney U-test all $p < 0.05$ after Bonferroni correction).

**F**. Quantification of the fraction of the total population of thalamic projections neurons responsive to glabrous skin (red) or hairy skin (blue) touch over developmental time. The percentage of neurons responsive to glabrous skin touch is stable (Kruskal-Wallis, $p > 0.05$), but the percentage of neurons responsive to hairy skin touch declines over developmental time (Kruskal-Wallis, Mann Whitney U-test all $p < 0.05$ after Bonferroni correction).

**Figure S5. LTMR optogenetic receptive field characterization**

**A**. A device for rapid, focal excitation of optogenetic proteins in skin. Laser pulses are passed through a beam expander and directed onto a set of galvometer scan mirrors. These mirrors are optically conjugate to the back aperature of a f-theta lens, such that rotation of the light beam translates a focal spot on skin. A high-pass dichroic allows images for the paw to be taken along the same optical path with an IR camera.

**B**. (left) X-mirror and Y-mirror command signal plotted above the power output of the laser measured with a calibrated photodiode. (right) An image of the laser spot illuminating the paw. The illuminated spot is ~50 microns in diameter.

**C**. Spatiotemporally complex optogenetic receptive fields are composed of isolated "RF spots." Spike-triggered average characterization of the responses to sparse white noise stimulation produce a receptive field description that give the probability of a spike at a location in the stimulus field at particular delay times from optical stimulation of the skin.

**D**. Similar to C, but in $Avil^{FlpO}$; $R26^{LsL\text{-}FsF\text{-}ReaChR::mCitrine}$ mice where Aβ LTMRs express ReaChR via injection of a AAV2/1 Cre virus into the dorsal column. These mice exhibit RF spots that are fewer in number but otherwise similar to (C), indicating that RF spots in (C) reflect multiple converging of Aβ LTMRs.

**E**. The number of spatiotemporally distinct RF spots in $Cdx2^{\text{Cre}}$ ; $Avil^{FlpO}$; $R26^{LsL\text{-}FsF\text{-}ReaChR::mCitrine}$ mice were all Aβ LTMRs express ReaChR and $Avil^{FlpO}$; $R26^{LsL\text{-}FsF\text{-}ReaChR::mCitrine}$ mice injected with AAV2/1 Cre into the dorsal column where ReaChR is expressed more sparsely.

**F**. The number of spikes evoked per stimulus to a RF spot in $Cdx2^{\text{Cre}}$ ; $Avil^{FlpO}$; $R26^{LsL\text{-}FsF\text{-}ReaChR::mCitrine}$ mice were all Aβ LTMRs express ReaChR and $Avil^{FlpO}$; $R26^{LsL\text{-}FsF\text{-}ReaChR::mCitrine}$ mice injected with AAV2/1 Cre into the dorsal column where ReaChR is expressed more sparsely.

**Figure S6. Genetic strategies for accessing glabrous and hairy Aβ-LTMRs that ascend the dorsal column**

**A**. The dorsal root ganglia of a $Ret^{\text{CreER}}$ , $R26^{LsL\text{-}FsF\text{-}tdTomato}$/ $R26^{LsL\text{-}F\text{-}ChR2\text{-}YFP\text{-}F}$ ; $Pvalb^{\text{Flpo}}$ mice. $\text{TdTomato}^{+}$ neurons are present at limb levels and are $\text{YFP}^{-}$, indicating that FlpO is expressed specifically in Aβ RA-LTMRs that innervate glabrous skin or Pacinian corpuscles and that this Cre-ON, FlpO-OFF strategy subtractively labels hairy skin Aβ RA-LTMRs.

**B**. Images of lanceolate endings in hairy skin and Meissner Corpuscle innervating neurons in glabrous skin from the genetic strategy in A, demonstrating genetic specificity.

**C**. Quantification of labeling efficiency in *Ret*$^{CreER}$ ; *Pvalb*$^{Flpo}$ ; *R26*$^{LsL\text{-}F\text{-}ChR2\text{-}YFP\text{-}F}$ mice. A significant fraction of NFH$^{+}$ lanceolate endings are labeled, while labeling was not detected in any Meissner Corpuscle innervating neurons examined (n=4 mice).

**D**. Validation of a genetic strategy that predominantly labels glabrous Aβ RA-LTMRs that project through the dorsal column of the spinal cord. We injected an AAV2-retro-tdTomato virus in the dorsal column of *Ntrk2*$^{CreER}$*; Avil*$^{FlpO}$*; R26*$^{LsL\text{-}FsF\text{-}ReaChR::mCitrine}$ mice and assessed co-localization of tdTomato with mCitrine.

**E**. Images of Meissner-associated TrkB+ Aβ LTMRs (top) and hairy skin lanceolate endings (bottom) from mice in (D).

**F**. Quantification of the percent of mCitrine$^{+}$ endings of each morphological type that are tdTom$^{+}$ and thus project to the dorsal column (n=2 mice).

**Figure S7. Piezo2 *in situ* hybridization analysis**

**A**. Sections of lumbar-level DRGs stained by *in situ* hybridization using probes against Piezo2 (exons 43-45) or Nefh, in *Cdx2-Cre; Piezo2*$^{flox/flox}$ animals or *Piezo2*$^{flox/flox}$ control siblings.

**B**. Quantification of the percentage of Nefh-positive cells that are Piezo2-positive. Three *Cdx2-Cre; Piezo2*$^{flox/flox}$ and two *Piezo2*$^{flox/flox}$ animals were quantified.

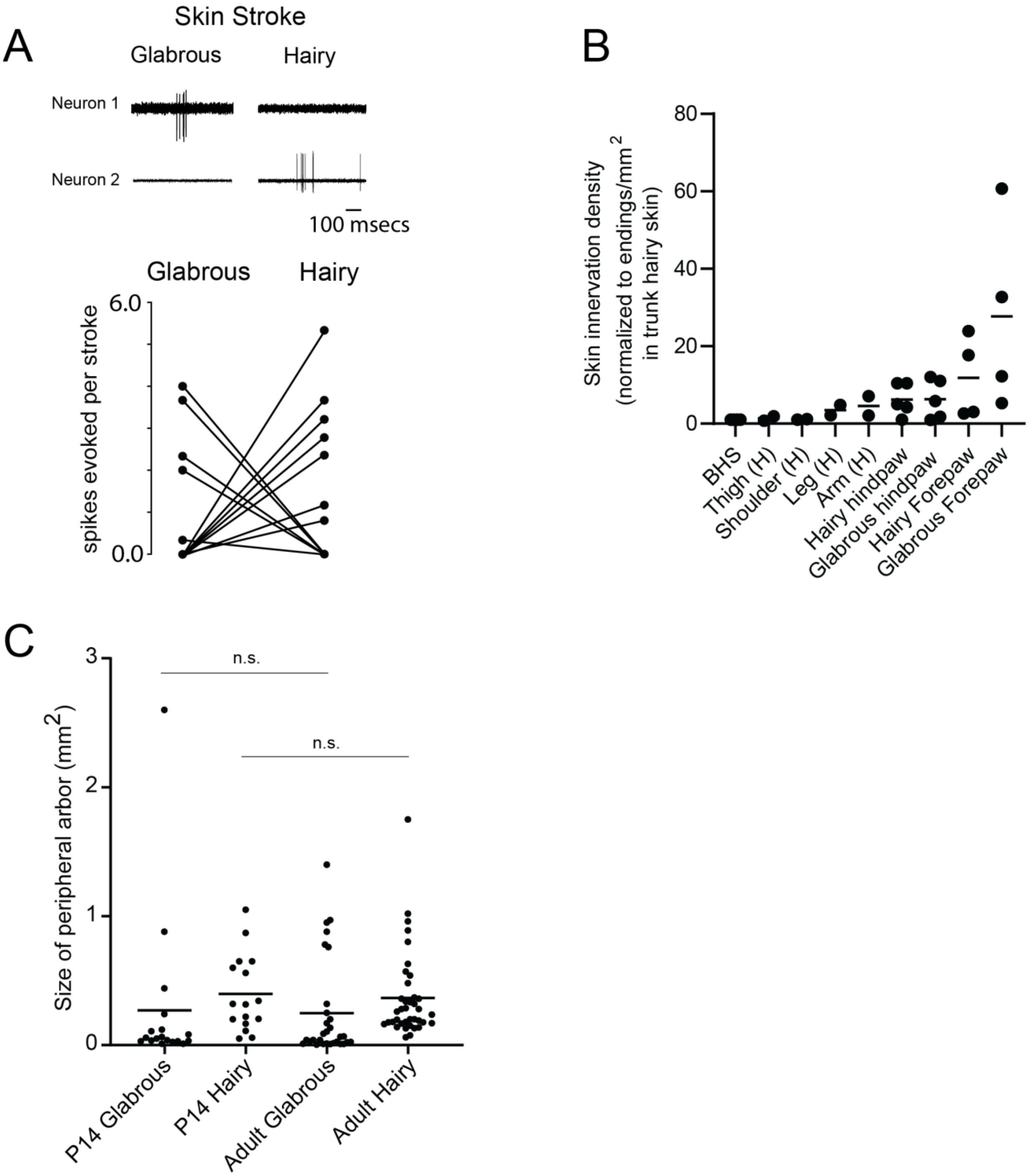


Supplemental Figure 1

### Three Distinct Populations of Neurons in the Gracile and Cuneate Nuclei

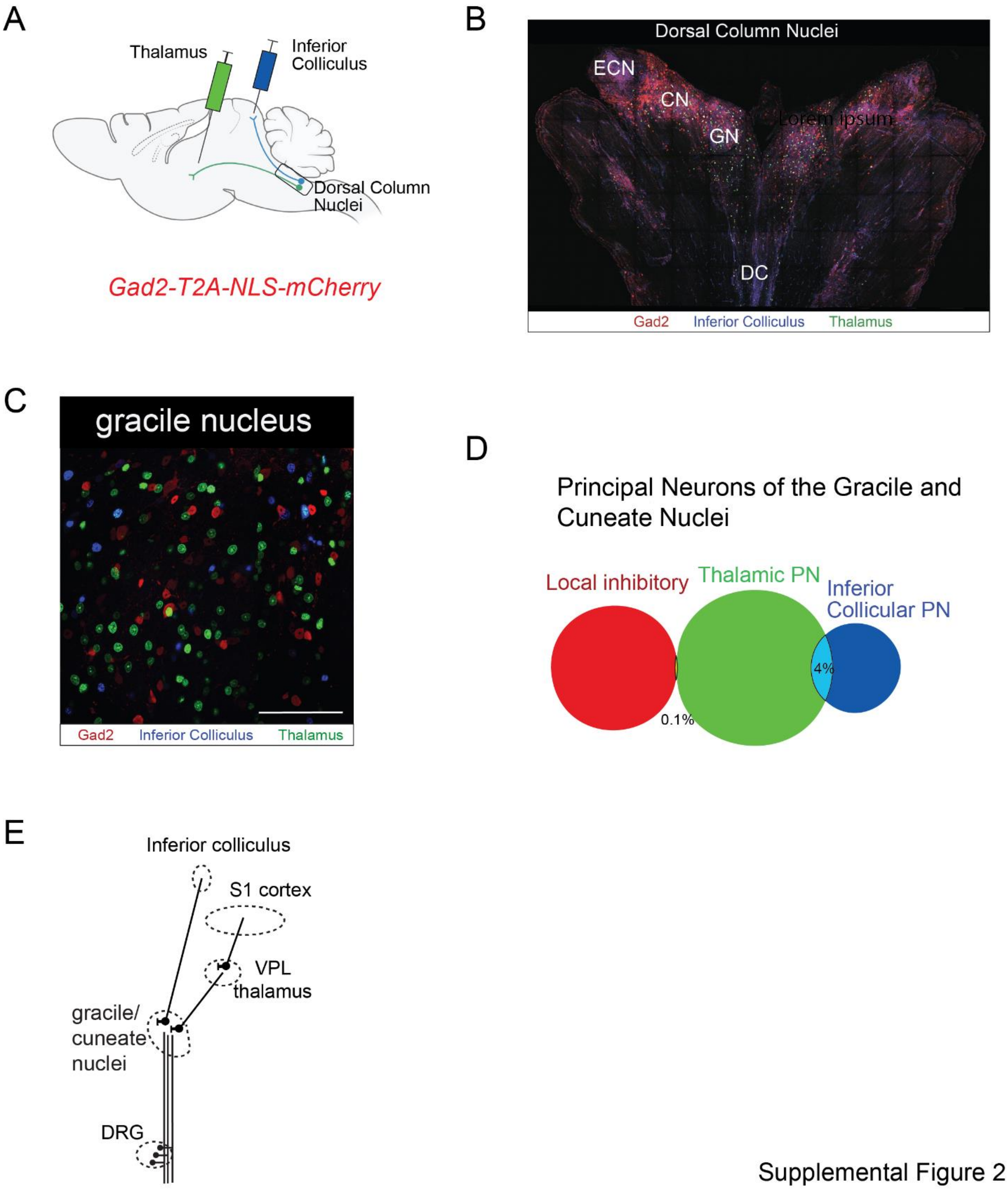


Supplemental Figure 2

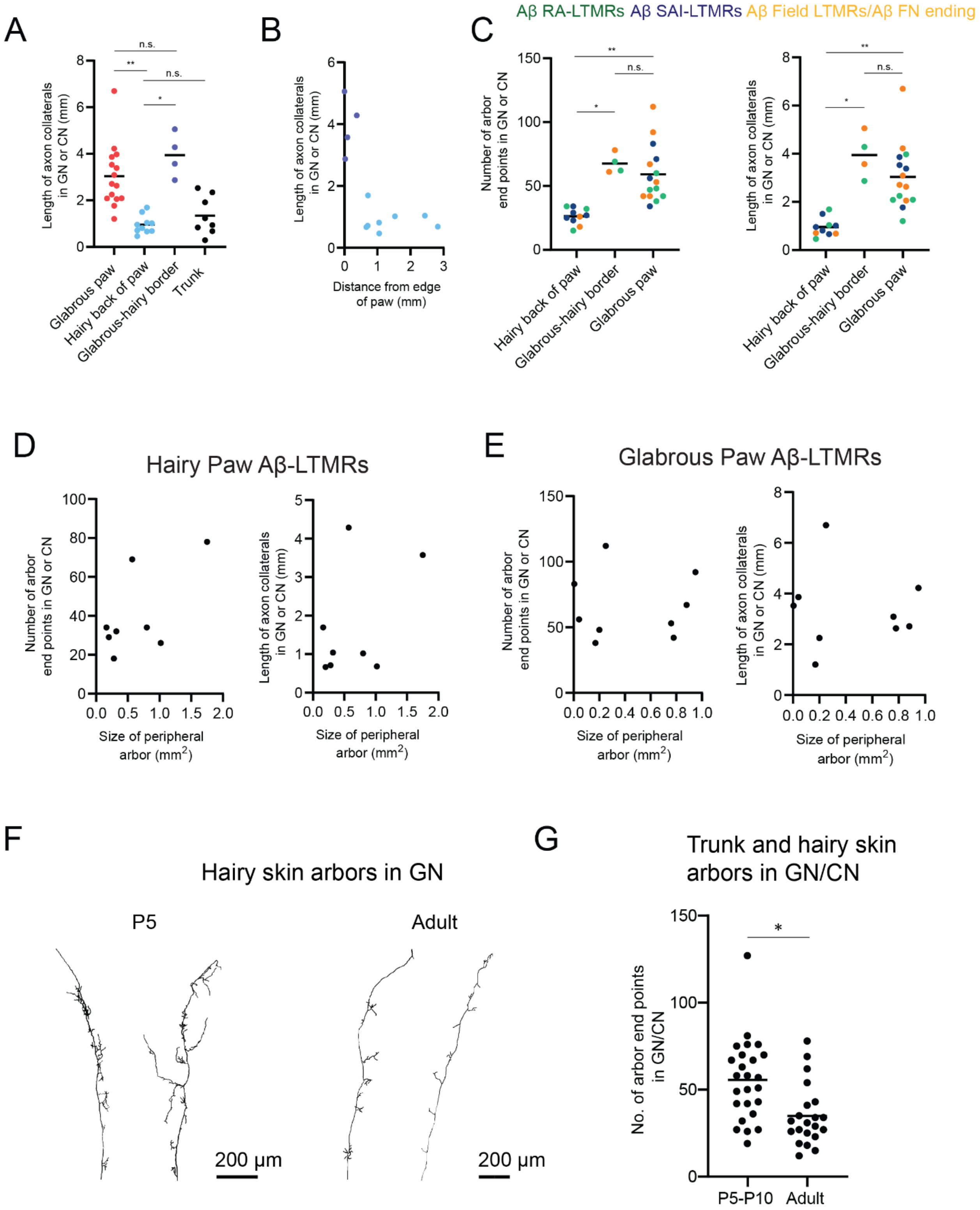


Supplemental Figure 3

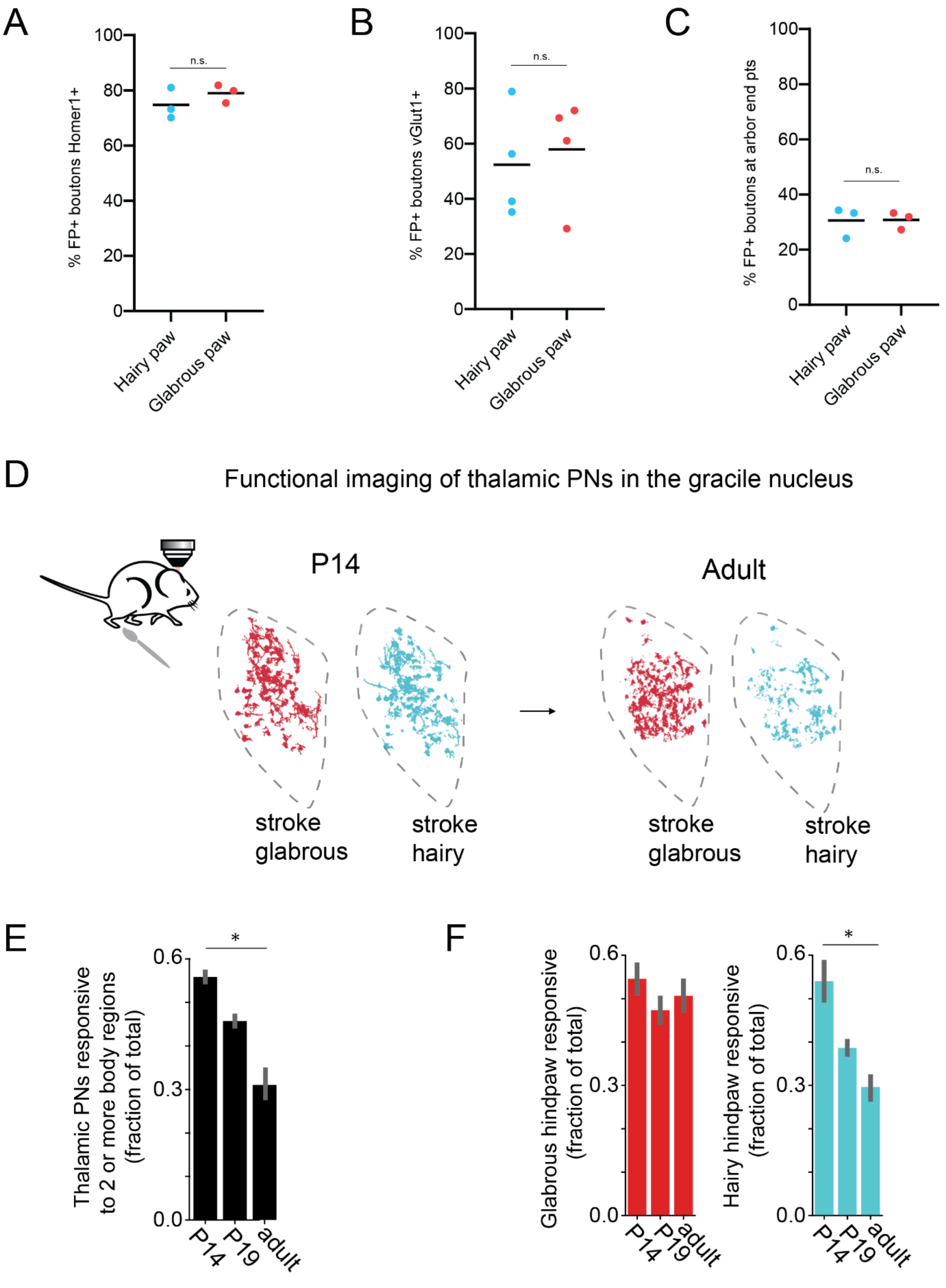


Supplemental Figure 4

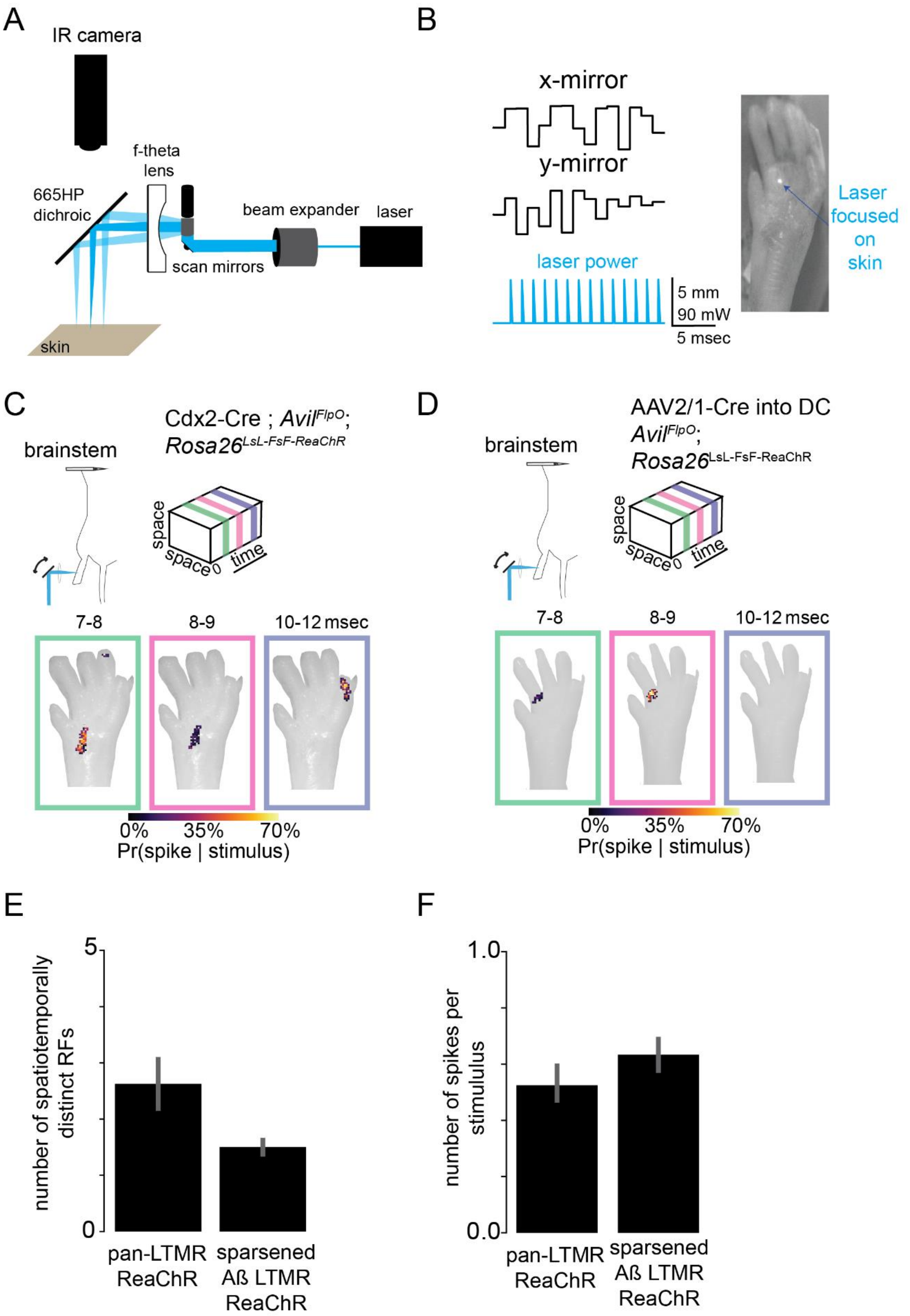


Supplemental Figure 5

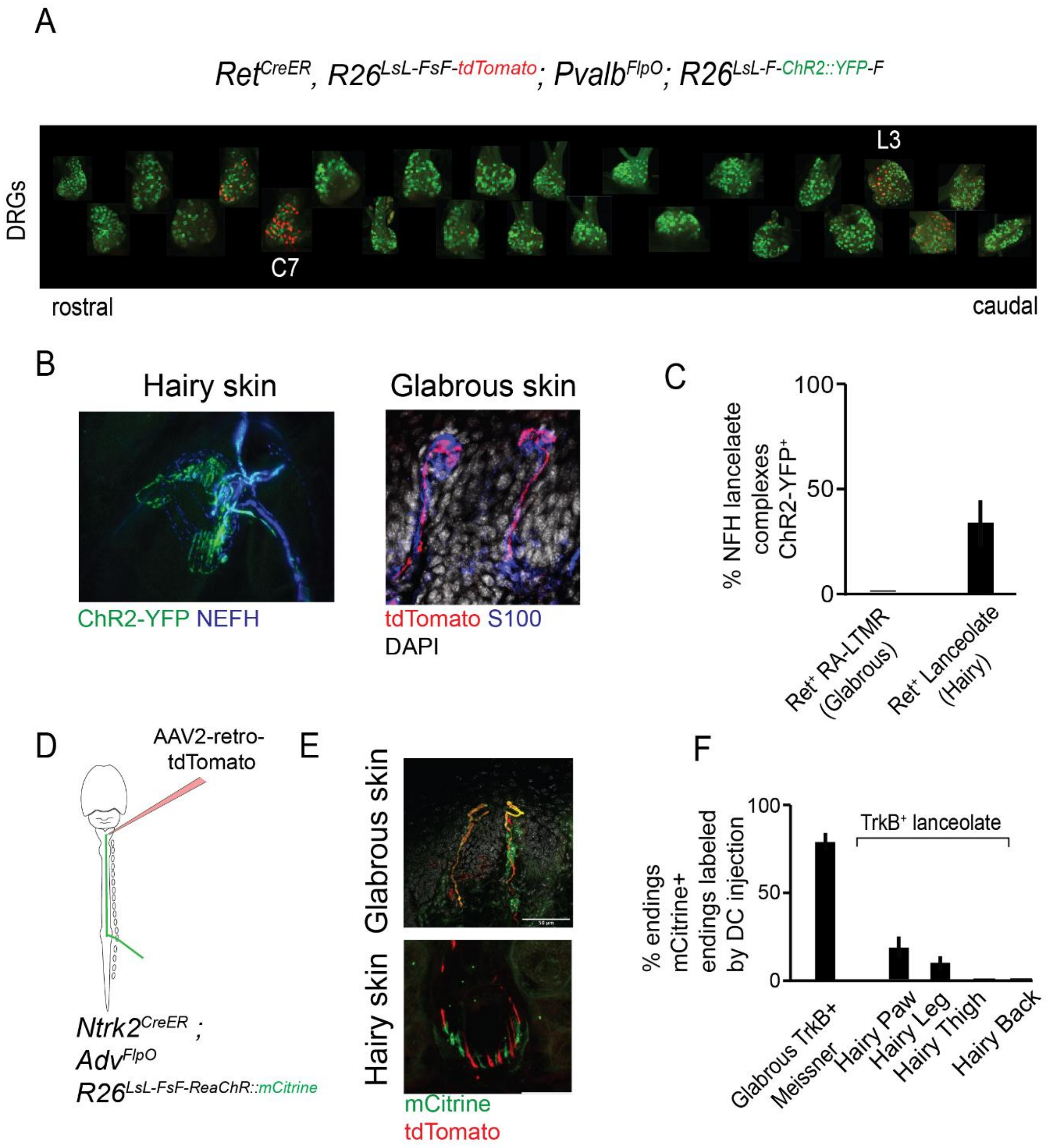


Supplemental Figure 6

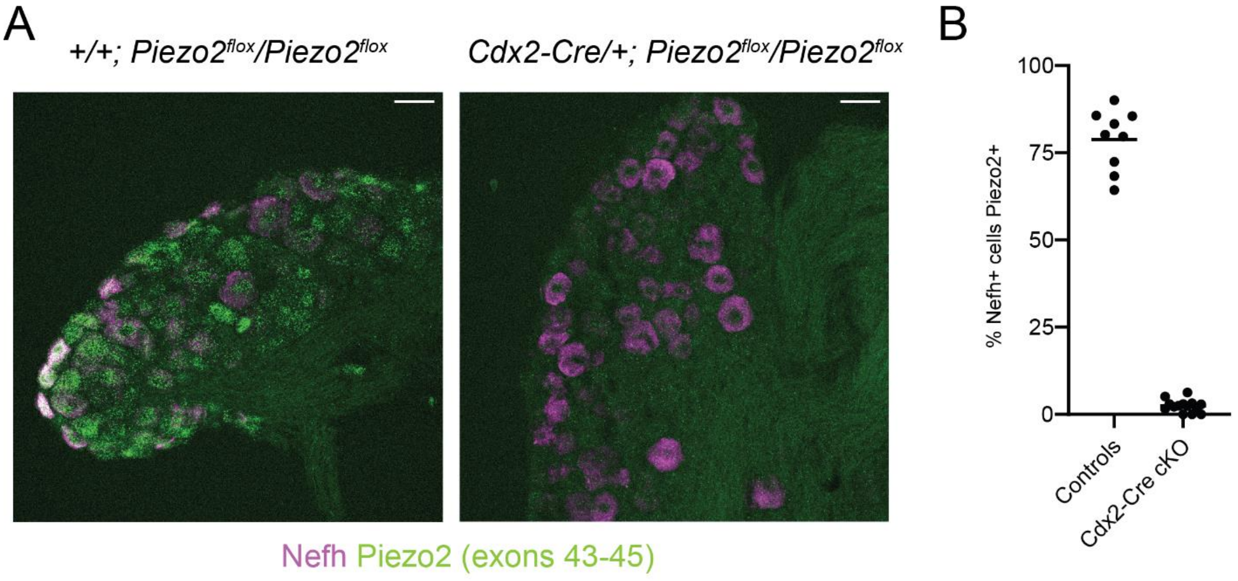


Supplemental Figure 7
